# A distinctive family of L,D-transpeptidases catalyzing L-Ala-mDAP crosslinks in Alpha and Betaproteobacteria

**DOI:** 10.1101/2023.10.31.564931

**Authors:** Akbar Espaillat, Laura Alvarez, Gabriel Torrens, Josy ter Beek, Vega Miguel-Ruano, Oihane Irazoki, Federico Gago, Juan A. Hermoso, Ronnie Per-Arne Berntsson, Felipe Cava

**Affiliations:** Department of Molecular Biology and Laboratory for Molecular Infection Medicine Sweden, Umeå Centre for Microbial Research, SciLifeLab, Umeå University, Umeå, Sweden; Department of Medical Biochemistry and Biophysics, Umeå University, Umeå, Sweden; Wallenberg Centre for Molecular Medicine, Umeå University, Umeå, Sweden; Department of Crystallography and Structural Biology, Institute of Physical Chemistry “Blas Cabrera”, CSIC, Madrid, Spain; Department of Biomedical Sciences & IQM-CSIC Associate Unit, School of Medicine and Health Sciences, University of Alcalá, E-28805 Madrid, Alcalá de Henares, Spain

## Abstract

Most bacteria are surrounded by an essential protective mesh-like structure called peptidoglycan, made of glycan chains crosslinked through short peptides by enzymes known as transpeptidases. Of these, penicillin-binding DD-transpeptidases connect adjacent peptide stems between their 4^th^ and 3^rd^ amino acids (4,3-type), D-alanine and a meso-diaminopimelic acid (mDAP) in Gram negatives, whereas LD-transpeptidases make the 3,3-type between mDAP^3^ residues. While these two processes explain the formation of crosslinks in most bacteria, recent investigations involving non-model species have brought to light novel crosslinking mechanisms that point to the existence of less-explored groups of peptidoglycan crosslinking enzymes. Here, we present the identification and characterization of a novel LD-transpeptidase found in the acetic acid bacterial *Gluconobacter oxydans*, named LDT_Go_, which performs 1,3-type crosslinks between L-Ala^1^ and mDAP^3^. LDT_Go_-like proteins are conserved among Alpha and Betaproteobacteria species that do not encode LD3,3-transpeptidases. Using a highly active ortholog, we demonstrated *in vitro* that this enzyme can work with non-terminal peptide bonds in the crosslinking process. This property is different from the strict specificity of typical LD- and DD-transpeptidases, which only deal with terminal peptide bonds. The high-resolution crystal structure of LDT_Go_ revealed significant distinctions when compared to 3,3-type LD-transpeptidases. These include a proline-rich region near the N-terminus that restricts substrate access to the active site, and an unprecedented cavity designed to accommodate both the glycan chain and the peptide stem from donor muropeptides, a feature that exhibits broad conservation among LD1,3-transpeptidases. Finally, we demonstrated the involvement of DD-crosslinking turnover in supplying the necessary substrate for LD1,3-transpeptidation. This phenomenon underscores the interplay between structurally distinct crosslinking mechanisms in maintaining cell wall integrity in *G. oxydans*.

## Introduction

Most bacteria are protected by an external cell wall that mainly consists of a net-like structure, named the peptidoglycan ^1^. Peptidoglycan, also known as murein *sacculus* (Latin word for “small bag”), functions as an exoskeleton that protects bacterial cells from bursting due to the elevated internal turgor pressure ^2^. Based on its organization, bacteria can be defined as Gram negative and Gram positives ^3^. Gram negative bacteria present a thin peptidoglycan monolayer confined in the periplasmic space, a cellular compartment between the cytoplasmic and the outer membranes. Gram positives have instead, a thick, multi-layered peptidoglycan outside of the cytoplasmic membrane and no outer membrane ^4^. At the compositional level, peptidoglycan is a heteropolymer made of glycan strands crosslinked by short peptides stems. The canonical peptidoglycan subunit (i.e., muropeptide) consists of the disaccharide pentapeptide N-acetylglucosamine (NAG) β(1→4) N-acetylmuramic acid (NAM)-L-alanine^1^-(γ)-D-glutamate^2^-(diamino acid^3^)-D-alanine^4^-D-alanine^5^ (abbreviated here as M5, for monomer disaccharide-pentapeptide). Usually, the diamino acid in the third position of the murein peptide stems is meso-diaminopimelic acid (mDAP^3^) in Gram negatives and L-Lys in Gram positives ^1^. However, various species display additional peptidoglycan chemical changes that typically enable adaptation of the cell wall to specific environmental challenges ^5^, e.g., substitution of the D-alanine at fifth position (D-Ala^5^) of the peptide moiety by D-lactate in vancomycin-resistant strains ^6^.

Peptidoglycan synthesis requires the coordinated action of synthetic and degradative enzymes that catalyze the insertion of new material into the pre-existing murein sacculus, thereby enabling sacculi expansion to support cell growth. Murein polymerization requires transglycosylases (TGases) and transpeptidases (TPases) activities. TGase enzymes include the bifunctional class A penicillin-binding proteins (PBPs), the shape, elongation, division, and sporulation proteins (SEDS), and the monofunctional glycosyl transferases ^7^. Peptidoglycan transpeptidation (i.e., crosslinking) is mainly catalyzed by class A and monofunctional class B PBPs, also called DD-TPases. However, many bacteria encode alternative TPases known as LD-TPases ^8^ that do not share sequence homology with PBPs. They present a YkuD-like domain (PFAM 03734) that includes a cysteine as the catalytic nucleophile instead of the conserved serine in PBPs ^8^. PBPs cleave the terminal peptide bond between the fourth and fifth amino acid of the donor pentapeptide (D-Ala^4^-D-Ala^5^) to form a new peptide bond connecting the D-Ala^4^ of the donor muropeptide with the D-chiral center of the mDAP^3^ from an adjacent acceptor muropeptide, thereby forming a 4,3-type or DD-crosslinking. Contrary to PBPs, LD-TPases primary cleave between mDAP^3^-D-Ala^4^ in the donor tetrapeptide and use the energy to form a crosslink between the L- and the D-center of two adjacent mDAP^3^ residues, thereby constituting a 3,3-type or LD-crosslink.

These are not the only differences between these two TPases. The activity of PBPs is vital for building a viable cell wall, and as a result, most bacteria carry one or multiple indispensable PBPs. Instead, LD-transpeptidation is not essential for survival, yet it is important for a number of processes such as peptidoglycan chemically editing by non-canonical D-amino acids (NCDAA) ^9^, the tethering of outer membrane proteins to the peptidoglycan ^10,11^, toxin secretion ^12^, lipopolysaccharide translocation ^13^, antibiotic resistance ^14^, and polar growth ^15^. We previously reported that the family Acetobacteriaceae presents a unique peptidoglycan chemical structure, which includes a novel type 1,3-type LD-crosslink between L-Ala^1^-DAP^3 16^. Using *Gluconobacter oxydans* as a model organism, we successfully identified the LD-TPase, i.e., LDT_Go_, responsible for the creation of these unconventional 1,3 peptidoglycan crosslinks. Following this, we conducted an in-depth exploration of its structural characteristics, biochemical properties, and its relevance in biological contexts. We show that LDT_Go_ has distinctive structural features with respect to 3,3 crosslinking enzymes, which includes the active site donor cavity and a N-proximal region that likely has a regulatory function with respect to its enzyme activity. Importantly, we demonstrate that contrary to what is known for other crosslinking enzymes, LD1,3-TPases have endopeptide-transferase activity, thus enabling these enzymes to use multiple donor muropeptides. We further show that although LDT_Go_ is constitutively expressed, LD1,3-crosslinking levels increase in stationary phase and we demonstrate this regulation depends on the DD-endopeptidase activity of PBP7, which controls substrate availability for LDT_Go_. This growth phase-dependent crosstalk between DD- and LD-crosslinking types supports peptidoglycan crosslinking homeostasis in this bacterium. Inactivation of LDT_Go_ sensitizes *G. oxydans* to cell envelope stresses, which cannot be complemented by canonical 3,3-type LD-crosslinks. Conservation of LDT_Go_-like enzymes within Alpha and Betaproteobacteria suggests an important role for this family of enzymes in shaping the cell wall of these organisms to adapt to challenging conditions.

## Results

### An uncharacterized YkuD-containing protein is involved in 1,3-type peptidoglycan crosslinking in *G. oxydans*

We previously found that the peptidoglycan of acetic acid bacteria (Acetobacterias) exhibits an unusual type of crosslink between the L-Ala at position 1 of the donor muropeptide stem peptide and the D-chiral center of mDAP at position 3 of the acceptor muropeptide (L-Ala^1^-DAP^3^) ^16^ (Fig 1A). We reasoned that an LD-TPase might catalyze this crosslinking reaction but found no homologs of the canonical LD-TPases that catalyze 3,3-crosslinks. As LD-TPases typically have a characteristic YkuD domain ^8^, we then searched for genes encoding YkuD-containing proteins in the genome of *G. oxydans* and found *gox2269* and *gox1074* (Fig. 1B). While the muropeptide profile of the clean deletion mutant strain Δ*gox2269* phenocopied the wild-type (Fig. S1), that of Δ*gox1074* exhibited a notable absence of 1,3-crosslinks, which were fully restored through ectopic complementation (Fig. 1C). This result demonstrates that Gox1074 plays an essential role in peptidoglycan LD1,3-crosslinking in *G. oxydans*.

**Figure 1.**
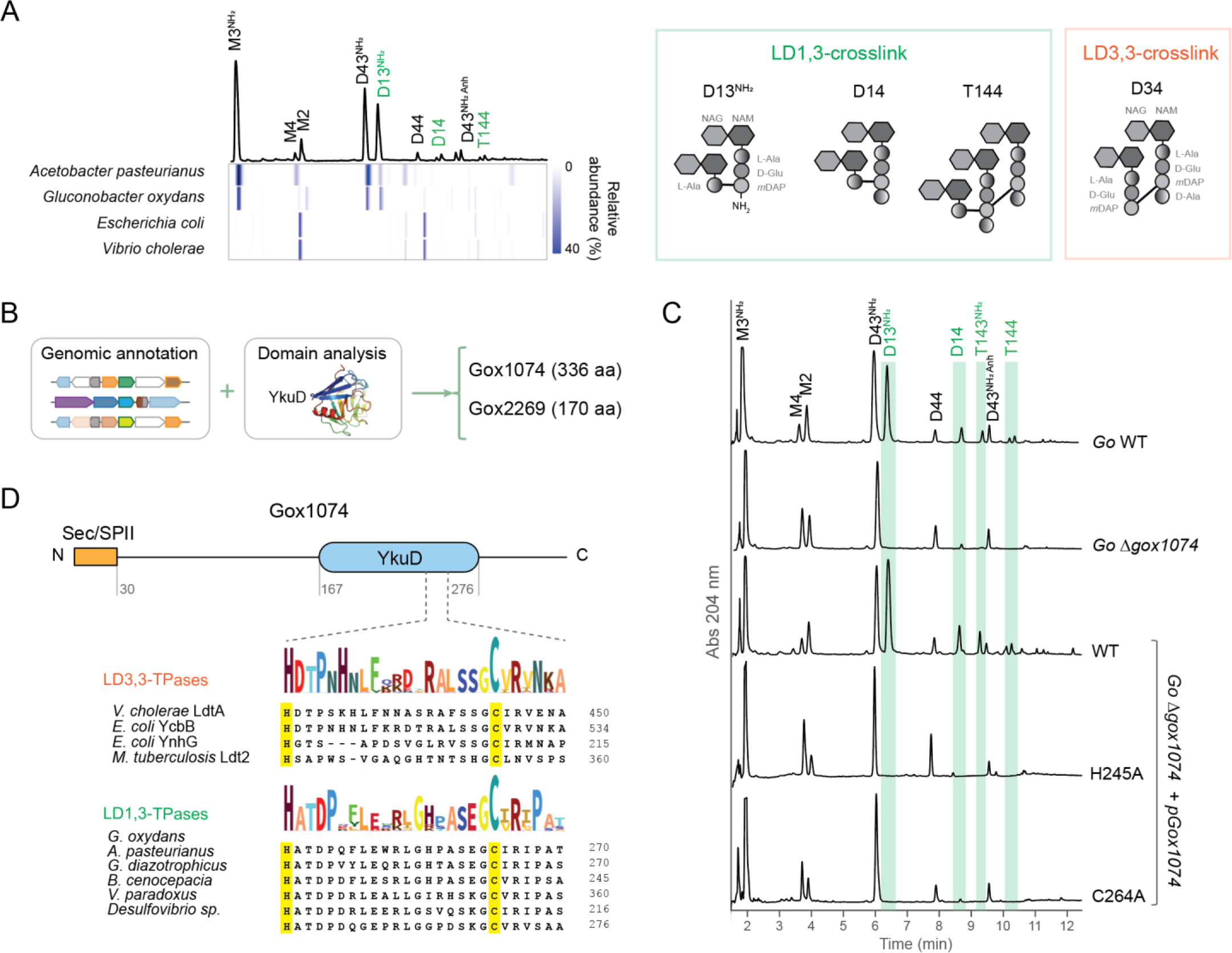
Identification of the enzyme catalyzing LD1,3-crosslink. A) Representative UV chromatogram of Gluconobacter oxydans peptidoglycan profile in stationary phase. LD1,3-crosslinked muropeptides are indicated in green (dimers, trimers and anhydro forms). Underneath, heatmap representing the relative abundance of each muropeptide in representative bacteria species from the Acetobacteriaceae family (Acetobacter pasteurianus and G. oxydans) and model organisms Escherichia coli and Vibrio cholerae. B) Scheme of the in silico search of YkuD-domain containing enzyme candidates responsible for LD1,3-crosslinking. C) Representative UV chromatograms of G. oxydans (Go) wild-type (WT), Δgox1074 and Δgox1074 pGox1074 complemented strains. LD1,3-crosslinked muropeptides are highlighted in green. D) Domain analysis of Gox1074. Details of the LDT conserved motif in the YkuD domain including the catalytic Cys and His residues highlighted in yellow.

The YkuD domain of Gox1074 includes the putative catalytic cysteine (C264) and histidine (H245) which are conserved throughout LD3,3-TPases (Fig. 1D). To assess if these residues are essential for Gox1074 function, we complemented the Δ*gox1074* strain with alleles in which these residues were replaced by alanine (C264A and H245A strains) and analyzed their peptidoglycan by UPLC-UV. Like Δ*gox1074,* the muropeptide profile of these strains completely lacked LD1,3-crosslinks, supporting that C264 and H245 are critical residues for Gox1074 activity (Fig. 1C).

SignalP 6.0 ^17^ predicted for Gox1074 an N-terminal Sec/SPII signal peptide typical of prokaryotic lipoproteins that includes the consensus sequence [LVI][ASTVI][GAS]C (Fig. S2A). By immunodetection, we demonstrated that Gox1074 is associated to the membrane protein fraction (Fig. S2B). For other TPases, it has been demonstrated that their association with the membrane facilitates critical protein-protein interactions for catalysis and localization ^18,19^. To investigate whether this is the case for Gox1074, we produced two additional Δ*gox1074* strains. In one instance, we ectopically expressed *gox1074* without its lipoprotein signal peptide (ΔSP) while in the other, we replaced it with the signal peptide from YcbB_Ec_ (SP_YcbB_), which is predicted to localize soluble in *E. coli*’s periplasmic space ^20^ (Fig. S2C). Our results suggest that while Gox1074 needs to be translocated to the periplasm to be functional, it does not require to be anchored to the membrane (Fig. S2DE). Based on these results we renamed Gox1074 as LDT_Go,_ for LDT of *G. oxydans*, and termed LDT_Go_-like enzymes as LD1,3-TPases to distinguish them from the LD-TPases performing 3,3 crosskings (LD3,3-TPases).

### Conservation of LD1,3-TPases

To analyze the conservation of LD1,3-TPases, we built a phylogenetic tree based on sequence similarity. We found that LDT_Go_ homologs are encoded in Alpha and Betaproteobacteria, particularly among the Acetobacteriaceae and Burkholderiaceae families but also in some Comamonadaceae, Alcaligenaceae and Oxalobacteraceae. Outside of these phyla, orthologs can also be found in some Desulfovibrionaceae species (Fig. 2A). Sequence alignment of representative LDT_Go_-like proteins belonging to the above indicated taxa showed these proteins conserve the catalytic YkuD domain but not the lipobox SP (Fig. 2B). Interestingly, LD1,3- and LD3,3-TPases appear to be mutually exclusive as homologs of LDT_Go_ and YcbB_Ec_ are not encoded in the same species (Fig. 2A).

**Figure 2.**
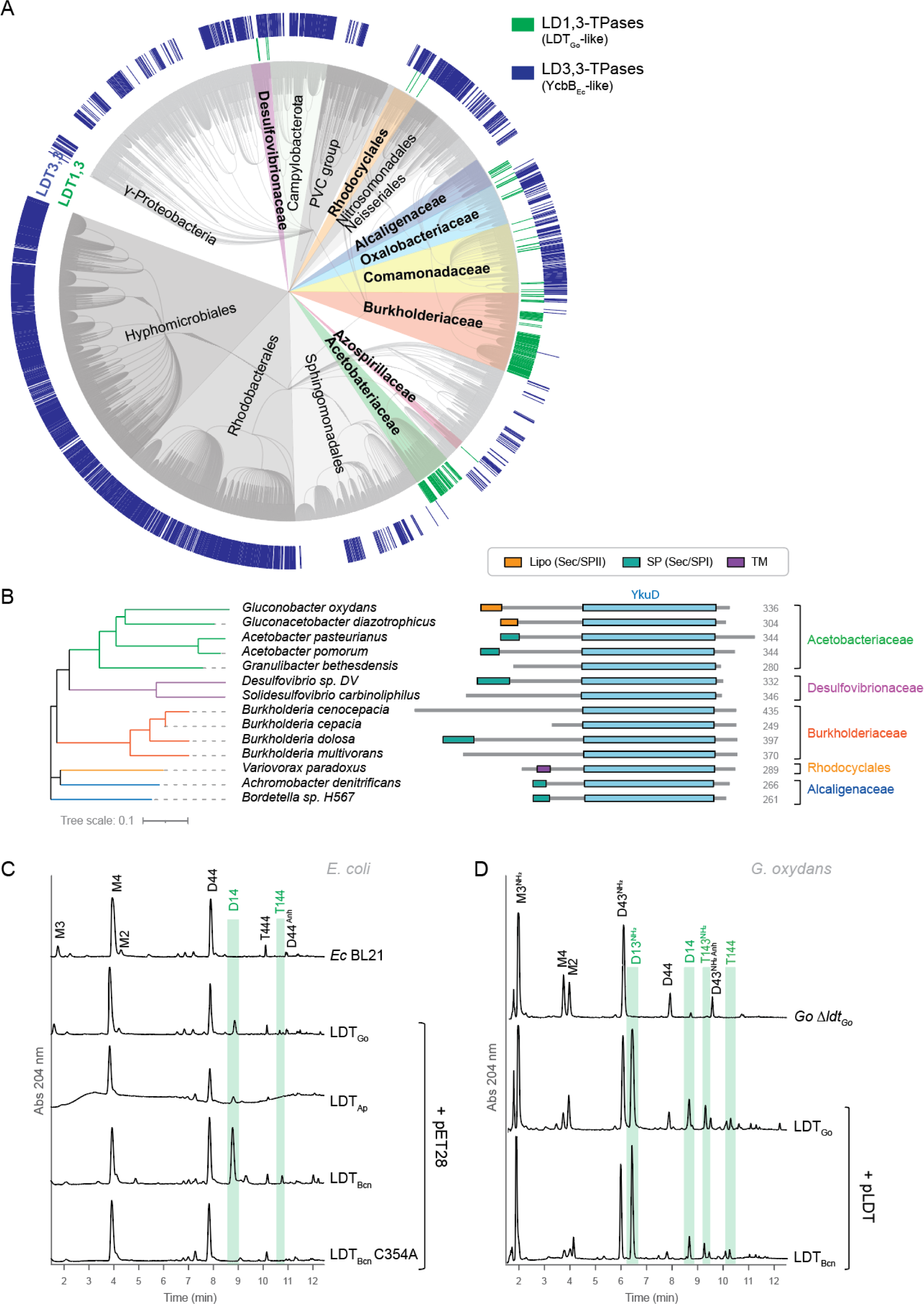
Conservation of LDT_Go_. A) Phylogenetic tree showing the conservation and distribution of LD-transpeptidases. In green, LDT_Go_-like proteins (LD1,3-TPases) and in blue, homologues to YcbB_Ec_ (LD3,3-TPases). Homology was assessed by BLAST against the NCBI and OrthoDBv11 databases. B) Comparison of the domains in representative LDT_Go_ orthologues from diverse bacterial Families. The presence of a signal peptide is indicated. Signal peptides and transmembrane domains are predicted using SignalP 6.0. C) UV muropeptide profiles of the heterologous expression of LDT_Go_ and its homologs from *Acetobacter pasteurianus* (LDT_Ap_), *Burkholderia cenocepacia* WT (LDT_Bcn_) and a catalytically inactive mutant (LDT_Bcn_ C354A) in *E. coli* BL21. LD1,3-crosslinked muropeptides are highlighted in green. D) UV muropeptide profiles of *G. oxydans* (*Go*) Δ*ldt_Go_* mutant and complemented derivatives expressing the LDT_Go_ and LDT_Bcn_. LD1,3-crosslinked muropeptides are highlighted in green.

To assess whether these predicted LDT_Go-_like proteins are authentic LD1,3-TPase enzymes, we expressed 2 orthologs in *E. coli* and subsequently analyzed the impact on the peptidoglycan. Both proteins produced LD1,3-crosslinks in *E. coli*, which is naturally devoid of these links, especially *Burkholderia cenocepacia* protein (referred to as LDT_Bcn_, BCAM2463), which generated much higher levels of the crosslink compared to both LDT_Go_ and its counterpart from *A. pasteurianus* (LDT_Ap_, NBRC3222_0766). As for LDT_Go_, LDT_Bcn_ activity was abolished by introducing a point mutation that changed the presumed catalytic cysteine to an alanine (C354A) (Fig. 2C) and successfully complemented *G. oxydans* Δ*ldt_Go_* (Fig. 2D). Interestingly, although expressing these enzymes in *E. coli* leads to substantial LD1,3-crosslinking, these muropeptide levels are comparatively scarce in their native species (Fig 1D, Fig. S3; ^16^), thereby suggesting the existence of species-specific regulatory mechanisms controlling the activity or expression of these enzymes.

Collectively, these findings indicate that LD1,3-TPase enzymes exhibit a high degree of conservation across various families within the Alpha and Betaproteobacteria that lack LD3,3-TPases.

### LD1,3-TPase donor substrate specificity

To make LD-crosslinked dimers, LD3,3-TPases use disaccharide tetrapeptides (M4) as donor muropeptide ^21^. By analogy, we reasoned that the disaccharide-dipeptide (M2) should serve as a donor in the production of LD1,3-crosslinked dimers. Nonetheless, this possibility appeared improbable due to the limited abundance of this muropeptide (5.4%) within the peptidoglycan of *E. coli*, particularly in contrast to the substantial production of the 1,3-crosslinked dimer D14 (20.5%) during the overexpression of LDT_Bcn_ in this bacterium (Fig. 2C). Given the significant reduction (34%) in M4 levels upon the expression of LDT_Bcn_, we postulated that M4, rather than M2, could serve as the main donor in LD1,3-transpeptidation reactions.

To investigate this hypothesis, we opted for LDT_Bcn_ over LDT_Go_ due to its superior activity in *E. coli* (Fig. 2C) and performed *in vitro* LD1,3-TPase assays on *V. cholerae* sacculi, which lacks LD1,3-crosslinks and only contains minimal levels of M2. We successfully detected D14 and trimers T144, confirming LDT_Bcn_ was active *in vitro* (Fig. 3AB, S4). As for LD3,3-TPases ^22^, synthesis of LD1,3-crosslinks were fully inhibited with imipenem and Cu^2+^ but not by ampicillin (Fig. 3C, S4). Notably, the formation of these LD1,3-crosslinked species was accompanied by a substantial decrease in M4, the main monomer, without a corresponding impact on M2 (Fig. 3AB). Additionally, we observed a significant reduction in D44 and the simultaneous generation of M43 (a disaccharide-tetrapeptide crosslinked to a tripeptide, Table S5) and M1, indicating that this enzyme also exhibits a notable endopeptidase activity *in vitro*, particularly on DD-crosslinked tetrapeptides (Fig. 3AB). These results further support our hypothesis that LDT_Bcn_ utilizes tetrapeptides as a substrate.

**Figure 3.**
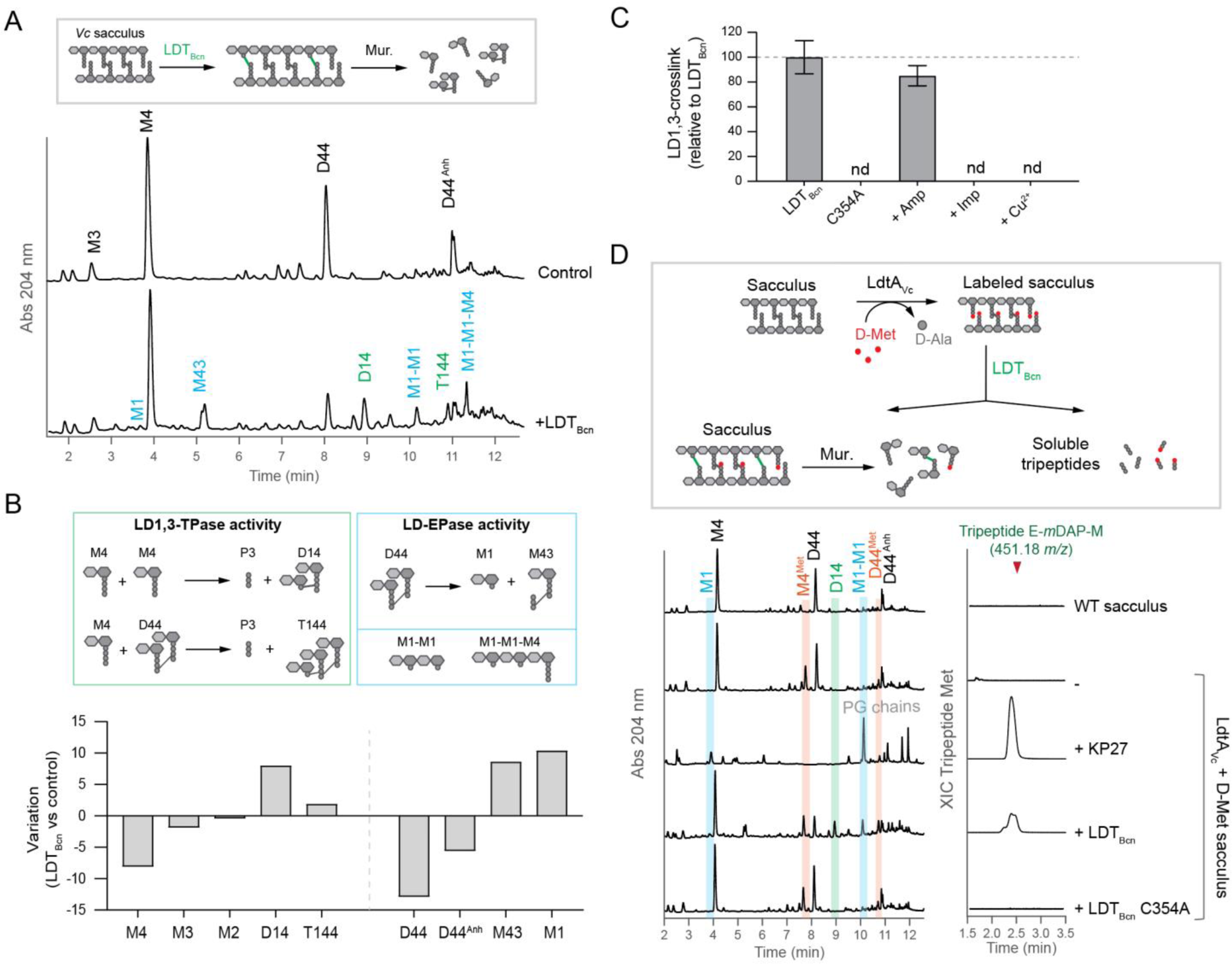
LD1,3-crosslinking activity. A) *In vitro* activity assays of LDT_Bcn_ on M4-rich peptidoglycan sacculi from *V. cholerae* vs control (no enzyme added). LD1,3-crosslinked muropeptides are labeled in green and the products of LDT_Bcn_ endopeptidase activity are labeled in blue. B) Muropeptide quantifications from panel A. Variation is calculated as the difference in relative molar abundance of the muropeptide in the LDT_Bcn_ *vs* control reaction in the *in vitro* assays. C) Effect of Ampicillin 100 µg/ml (Amp), Imipenem 100 µg/ml (Imp) and copper 1 mM (Cu^2+^) on the *in vitro* assays shown in panel A. nd: not detected. D) Scheme and *in vitro* activity assays of LDT_Bcn_ on M4-rich peptidoglycan sacculi previously modified with incorporated D-Met at the terminal position of the tetrapeptides. KP27 endopeptidase cleaving between L-Ala^1^ and D-Glu^2^ was used as control. Left side, UV muropeptide profiles of the muramidase-digested insoluble pellets showing the presence of D-Met modified muropeptides (in red), endopeptidase products (M1 and M1M1, in blue) and LD1,3-crosslinked muropeptides (in green). The MS extracted ion current (XIC) trace of the D-Met containing tripeptide (E-*m*DAP-M) in the soluble fraction is shown on the right side.

We then reasoned that if LDT_Bcn_ uses M4 as a donor muropeptide, we should observe the release of the tripeptide D-Glu-mDAP-D-Ala when D14 is generated (Fig. 3B). To investigate this hypothesis, we used the phage endopeptidase KP27 ^23^ as control, which cleaves peptidoglycan between L-Ala^1^-D-Glu^2^ and thus releases the same tripeptide (Fig. S5A). However, even though KP27 produces high levels of M1 muropeptide and significantly reduces M4 levels, we were unable to detect D-Glu-mDAP-D-Ala, likely because it elutes at the solvent front (Fig. S5B). To address this challenge, we initially substituted the terminal D-Ala^4^ by D-Met, using LdtA ^9^ (Figure 3D). We considered that the delayed elution of the more hydrophobic D-Met-containing tripeptide (D-Glu-mDAP-D-Met) would aid in its detection (Figure 3D). Indeed, this time we successfully identified both M1 and D-Glu-mDAP-D-Met in the peptidoglycan samples treated with KP27 (Fig. 3D). Remarkably, when D-Met-labeled peptidoglycan was treated with LDT_Bcn_, it generated D14 dimers and M1 muropeptides, and released the D-Glu-mDAP-D-Met tripeptide, thereby confirming this LD1,3-TPase can use M4 as the donor muropeptide in transpeptidase and endopeptidase reactions (Fig. 3D).

To explore whether M3, like M4, can also serve as the donor muropeptide, we conducted comparable *in vitro* LD1,3-transpeptidation assays. However, this time, we applied these assays to sacculi in which the monomer tetrapeptides were enzymatically trimmed to tripeptides by an LD-carboxypeptidase (Fig. S6A) ^24^. As expected, LDT_Bcn_ activity on this “M3-enriched” sacculi produced LD-crosslinked dimers (D13), M43 and M1 levels that were on par with those measured when using “M4-rich” peptidoglycan as substrate (Fig. S6BC). Once more, no changes in the M2 levels were observed between the samples subjected to LDT_Bcn_ treatment and those left untreated reinforcing the notion that M2 is not the favored donor muropeptide (Fig. S6BC). Interestingly, no LD1,3-crosslinked muropeptides were detected when we used a muramidase-digested peptidoglycan or purified muropeptides. This observation suggests a preference of the enzyme for peptidoglycan chains rather than individual muropeptides.

In summary, these results conclusively establish that LD1,3-TPases represent a distinct category of endopeptide-transferases which conduct L-Ala^1^ and D-Glu^2^ endopeptidation on donor muropeptides with diverse peptide stem lengths, ultimately enabling the formation of LD1,3-crosslinks.

### LD1,3-TPase perform D,L-amino acid exchange reactions

Additional differences in the catalysis between LD1,3- and LD3,3-TPases were observed when studying their ability to perform amino acid exchange reactions. As LD3,3-TPases can exchange the D-Ala^4^ of the peptide stems by non-canonical D-amino acids (NCDAA), fluorescent D-amino acids (FDAA) or clickable DAA ^9,25,26^, we wondered whether LD1,3-TPases could also perform a similar D,D-amino acid exchange reaction (Fig. 4A). However, neither incorporation of D-Met or the FDAA HADA was detected in *G. oxydans* wild-type cells compared to a derived strain in which the *ldt_Go_* locus was chromosomally replaced by *ycbB_Ec_* (Δ*ldt_Go_*::*ycbB*) (Fig. 4BC, S7). Interestingly, by UPLC-MS we detected trace amounts of several M2 ions in the peptidoglycan of *G. oxydans* that were absent in the *Δldt_Go_* mutant strain, corresponding to dipeptides that had exchanged the D-Glu^2^ by Phe and Trp (Fig. 4D, S8 and S9). These M2^Phe^ and M2^Trp^ species were present also in *E. coli* overexpressing the LDT_Go_ wild-type but not its catalytically inactive C264A mutant version (Fig. S9B). As neither *G. oxydans* nor *E. coli* encode homologs to the broad-spectrum racemase BsrV ^27,28^, the enzymes that produced NCDAAs, we reasoned that Phe and Trp were likely L-amino acids. To test this hypothesis, we first treated *G. oxydans* sacculi (which contain significant levels M2) with LDT_Bcn_ *in vitro* in the presence and absence of L-Phe and D-Phe. However, we found no significant changes in the M2^Phe^ levels between these samples, suggesting this activity might not be detectable *in vitro* (Fig. S10). Therefore, we grew *G. oxydans* wild-type and *Δldt_Go_* strains in cultures supplemented with 10 mM of L-Phe or D-Phe and monitored the generation of the M2^Phe^ muropeptide. Our results detected increased levels of M2^Phe^ (2.5-fold) only in the wild-type cultures supplemented with L-Phe, suggesting that in addition to its LD1,3-crosslinking activity, LDT_Go_ has D,L-amino acid exchange activity (Fig. 4E).

**Figure 4.**
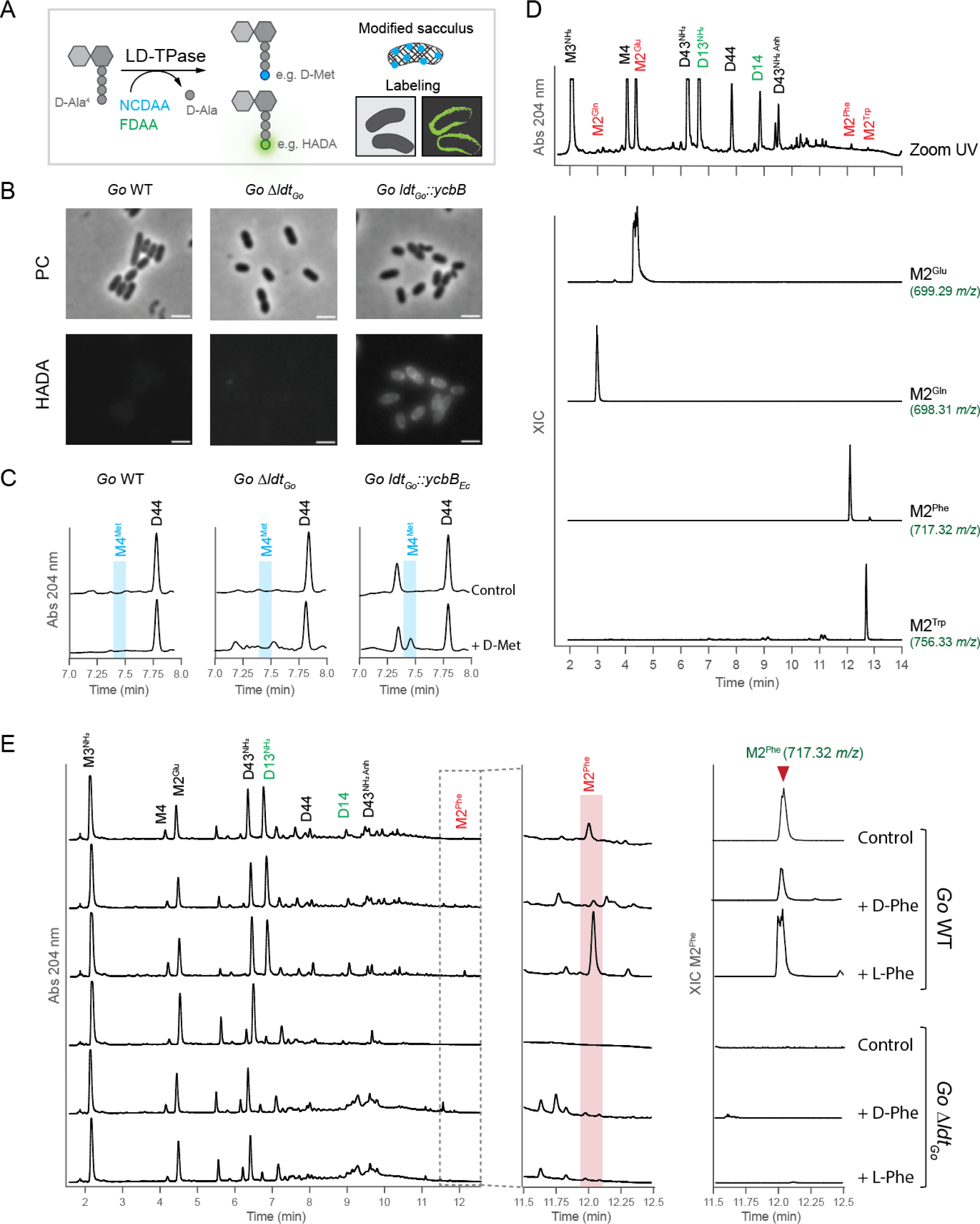
LDT_Go_ catalyzes D,L-amino acid exchange reaction. A) Scheme of the amino acid exchange reaction performed by LD3,3-TPases with non-canonical (NCDAA. e.g., D-Met) and Fluorescent (FDAA, e.g., HADA) D-amino acids. B) Phase contrast (PC) and fluorescence microscopy of *G. oxydans* wild-type (WT), Δ*ldt_Go_* mutant and *ldt_Go_::ycbB_Ec_* allelic exchange cells labeled with HADA. Scale bar: 2 µm. C) Zoom-in of the UV muropeptide profiles of the same strains indicated in panel B cultured with 10 mM of D-Met or without (control). D) Zoom-in of the UV muropeptide profile of *G. oxydans* WT highlighting the dipeptide muropeptides (M2^X^, in red) including those exhibiting amino acid exchange (M2^Phe^ and M2^Trp^), and the LD1,3-crosslinked muropeptides in green. The MS extracted ion current (XIC) traces are shown for the indicated M2^X^ muropeptide species. E) UV muropeptide profile of *G. oxydans* (*Go*) WT and Δ*ldt_Go_* mutant strains cultured without (control) or with 10mM of L-Phe or D-Phe (left panel). Middle panel: zoom-in of the UV muropeptide profile where the M2^Phe^ muropeptide elutes. Right panel: MS extracted ion current (XIC) trace of M2^Phe^.

### Three-dimensional structure of LDT_Go_

To understand the distinctive structural and catalytic characteristics of LD1,3-TPase enzymes in relation to their LD3,3-TPase counterparts, we successfully determined the crystal structure of a functional soluble LDT_Go_ variant at a resolution of 1.7 Å (PDB ID: 8QZG, Fig. 5A, Fig. S11). The electron density allowed us to build a model for residues 54 to 331, except for residues 201 to 215 corresponding to the capping loop that are likely in a flexible region. Comparison of the structures of LDT_Go_ and YcbB_Ec_ (PDB ID: 6NTW), revealed that while both proteins present YkuD domains with superimposed catalytic Cys and His residues, the overall structure of the domain is rather different (Fig. 5B). One striking change is in the capping loop, which is larger in YcbB_Ec_ and smaller and partially disordered in LDT_Go_. A second major difference is in the size and conformation of the β sheet and loops surrounding the catalytic center (Fig. 5B). As detailed below, these structural differences are essential to understand the unique catalytic properties exhibited by LDT_Go_.

**Figure 5.**
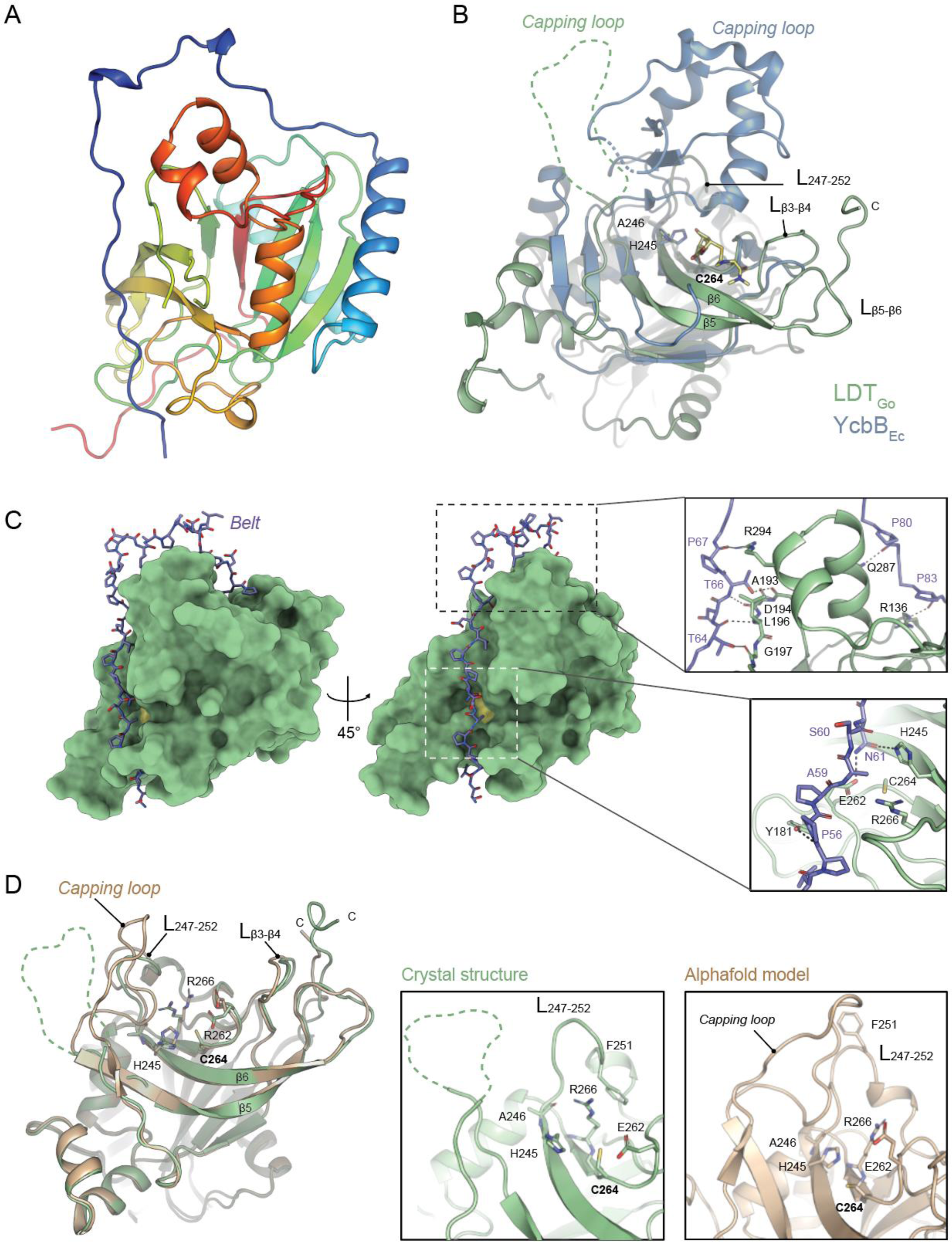
Structure of LDT_Go_. A) Overview of the LDT_Go_ structure, colored from the N-terminus (blue) to the C-terminus (red). B) Superimposition of LDT_Go_ (green) and the catalytic domain of YcbB_Ec_ (blue) bound to meropenem (yellow). Three characteristic features are highlighted, i) the capping loop of both enzymes, which in LDT_Go_ is disordered, ii) the LDT_Go_ catalytic residues C264 and H245, and iii) the distinctive LDT_Go_ domain within the active site composed of 2 interconnected loops between the β-strands 3 and 4 (L_β3-β4_) and the β-strands 5 and 6 and (L_β5-β6_). C) Surface representation of LDT _Go_ (green) with the Pro-rich belt shown as sticks (C-atoms colored in purple). The catalytic Cys is shown in yellow. Zoom-in panels show the hydrogen bonds formed between the belt and the rest of the protein. D) Superimposition of the LDT_Go_ structure (green) and its Alphafold2 prediction model (light brown), with the main differences highlighted in the zoom-in panels.

LDT_Go_ contains an elongated and unstructured proline-rich N-proximal region, spanning until residue 86, which we will refer to as the “belt” for reasons we explain below. Although this region demonstrates variability in its amino acid sequence, it is broadly conserved among LD1,3-TPases, while being conspicuously absent in LD3,3-TPases (Fig S12). In our crystal structure, the belt wraps around the protein, forming up to 11 hydrogen bonds, various van der Waals interactions and inserts into the catalytic cavity of the YkuD domain (Fig. 5C, Table S1). It is noteworthy that Asn61 from the belt is making hydrogen bonds with two residues located at both sides of the catalytic Cys264 (Fig.5C), the catalytic His245 and the Glu262 that, as described below, is part of the acidic patch close to the acyl-donor site. Thus, the obtained crystal structure of LDT_Go_ likely represents a self-inhibited state in which the belt occludes the active site from the peptidoglycan and blocks essential catalytic residues.

In the crystal structure, the capping loop is disordered and the L_247-252_ presents a conformation in which the Phe251 traps Arg266 from the active site through a cation-π interaction (Fig. 5D, middle panel). We ran molecular dynamics (MD) simulation with the crystal structure showing that the belt and the capping loop present the larger variations along the 350 ns simulation (Fig. S13A and Movie S1). Furthermore, MD simulations in which we removed the belt *in silico* generated a model that was remarkably similar to that predicted by AlphaFold2 (AF) for LDT_Go,_ where the belt does not encircle the protein, allowing the active site to remain unobstructed (Fig. S13B). In this model, the capping loop approaches the active site, the L_247-252_ partially refolds and thus Phe251 moves away from the active site. This allows a reorientation of Arg268 that can then make a salt-bridge interaction with Glu262 (that in the crystal structure makes a H-bond with Asn61 from the belt) (Fig. 5).

In summary, two different conformations are observed for LDT_Go_. One of these conformations involves a self-inhibited state, as observed in our crystal structure. In this state, the belt affects the conformation of catalytic residues and the loops around the active center thereby impeding access to the muropeptide substrates. The second conformation, represented by MD simulations and the AF model, likely features an active state of the protein. Here, the belt has moved away, and both the capping loop and the L_247-252_ region have refolded to expose the substrate-binding site. We subsequently leverage this model of the active protein to investigate the active site of LDT_Go_ and to shed light on the mechanisms by which LD1,3- and LD3,3-TPase enzymes catalyze their specific crosslinks.

### Structural basis of the 1,3 transpeptidation activity of LDT_Go_

As detailed before, our *in vivo* and *in vitro* experiments demonstrate that LDT_Go_ performs LD1,3-crosslinks by transferring the energy of a non-terminal peptide bond (e.g., using M4 as donor muropeptide). We reasoned that this unprecedented transpeptidation activity should be explained by specific structural modifications in the active site of LD1,3-TPases. We thus performed an in-depth analysis of the LDT_Go_ active site compared to that of the LD3,3-TPase YcbB_Ec_ in complex with meropenem (PDB ID: 6NTW) (Fig. 6A-D, S14).

**Figure 6.**
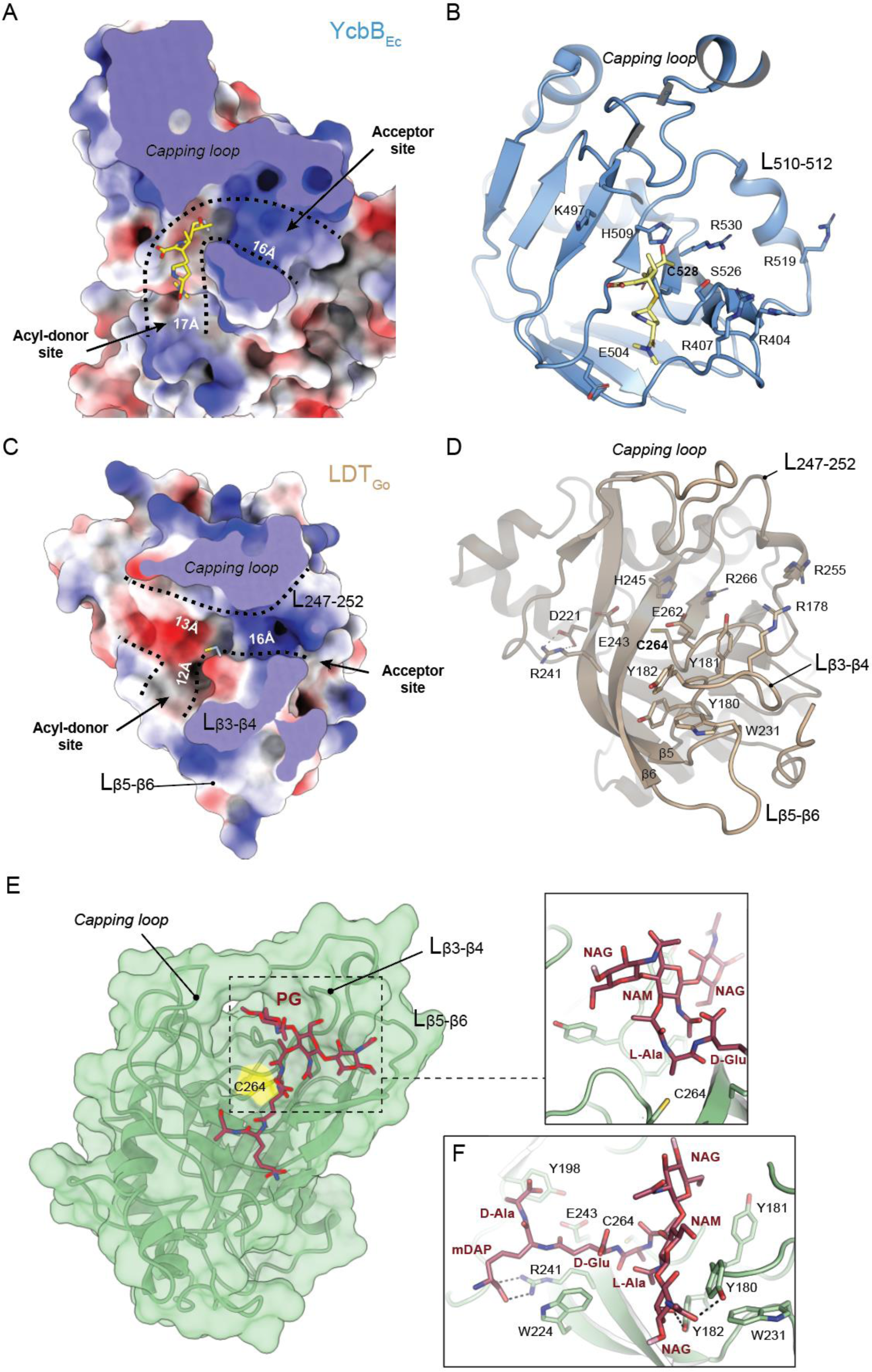
LDT_Go_ active center. Electrostatic surface representation, ranging from –13.35 to +15.88 for YcbB_Ec_ (A), and from –15.69 to +12.77 for LDT_Go_ (C) and ribbon diagram (B, D) of the donor and acceptor sites of YcbB_Ec_ in complex with the antibiotic meropenem (in yellow) (A, B) and of LDT_Go_ (C, D). The dimensions of the donor and acceptor cavities as well as the positions of the relevant loops and residues are indicated. E) LDT_Go_ with the disaccharide tetrapeptide (M4) modelled in the active site. Molecular surface of LDT _Go_ (green) with the catalytic C264 highlighted in yellow. The muropeptide is represented as sticks (C atoms colored as dark red). The positions of the relevant loops are indicated by arrows and labeled. F) Detailed view of the interaction of the M4 muropeptide into the LDT_Go_ active site. Relevant residues in the protein are represented as sticks and labeled.

In addition to the previously mentioned differences in the capping loop size (smaller in LDT_Go_), we observed very important differences in some of the β strands that define the active site in LDT_Go_. In particular, β5 and β6 (in which the catalytic His245 lies) exhibit significantly greater length in LDT_Go_ compared to YcbB_Ec_. Further, these two β strands are linked by the extended loop L_β5-β6_, a structural feature that is absent in YcbB_Ec_ (Fig. 6B, 6D, S14).

Another distinctive feature of LDT_Go_ is the presence of a long loop connecting β3 and β4, L_β3-β4_, that includes a unique triad of *in-tandem* Tyr residues (Tyr180, Tyr181 and Tyr182) (Figs. 6D, S14, S15). The side-chain orientation of two of these Tyr residues is stabilized through the interaction with Trp231 from the L_β5-β6_ (Fig. 6D). Interestingly, this aromatic triad together with the associated Trp is broadly conserved among LDT_Go_ orthologs found in Acetobacteriaceae species (Fig. S15AB). Structural predictions by AF of orthologues from different phyla underscores that the above-mentioned distinctive features of LDT_Go_ namely, the longer β5 and β6, and loops L_β5-β6_ and L_β3-β4_, are structural landmarks of this new family of L,D-TPases (Fig. S15C-I).

All these modifications together with a different distribution of acidic and basic residues likely account for the crosslinking specificity of LD1,3-TPases. While the acceptor groove exhibits similar basic characteristics and dimensions in both YcbB_Ec_ and LDT_Go_ (Fig. 6AC), it is the donor site that stands out as notably distinct in these proteins. In LDT_Go_, the donor site, marked by the position of meropenem in the YcbB_Ec_:meropenem complex, is significantly shorter, measuring around 12 Å compared to the longer span of about 17 Å in YcbB_Ec_. Additionally, in YcbB_Ec_, there is a narrow groove that extends toward the catalytic Cys residue, whereas in LDT_Go_, a large and open cavity is observed near the donor site with a pronounced acidic character (Fig. 6C). MD simulations confirmed that the LDT_Go_ donor site can stabilize M4 to make LD1,3-crosslinked dimers. Our results with a tetrapeptide ligand connected to a short (Fig. 6E) or longer (Fig. S16) peptidoglycan glycan chain, indicate that the capping loop and L_β5-β6_ and L_β3-β4_ are crucial elements in the stabilization of glycan chains of the donor substrate. In particular, the triad of Tyr residues (Tyr180, Tyr181 and Tyr182) in loop L_β3-β4_ and the Trp231 in L_β5-β6_ are involved in both van der Waals and H-bond interactions with the three sugar rings (Fig. 6F). This characteristic sets LDT_Go_ apart from LD3,3-TPases, which primarily target the peptide stem ^29^. This structural aspect facilitates the approach of the NAM component of the glycan moiety of the muropeptide towards the catalytic Cys264, ultimately enabling the formation of an adduct with L-Ala^1^ (Fig. 6E). Remarkably, the remaining part of the peptide moiety (i.e., the tripeptide D-Glu^2^-mDAP^3^-D-Ala^4^) is located in the extra cavity, not present in LD3,3-TPases, by the donor groove of LDT_Go_. The tripeptide after the L-Ala^1^ is stabilized by van der Waals interactions (mainly performed by Trp224 and Tyr198) and only a strong polar (salt-bridge) interactions is predicted to occur between carboxylate group of mDAP and Arg241 (Fig. 6F). The acidic character observed in this extra region could provide a repulsive effect to avoid the carboxylate-containing residues at positions 2-4 (i.e., D-Glu-M-DAP-D-Ala) to be placed close to the catalytic Cys residue. Thus, our MD simulations thereby support the preferential utilization of M3 and M4 muropeptides and explain the unique features observed in LDT_Go_ compared to LD3,3-TPases.

### PBP7 endopeptidase regulates LD1,3-crosslinking in *G. oxydans* by restricting the supply of peptidoglycan tetrapeptide monomers

LD1,3-crosslinks are more prominent in the stationary phase peptidoglycan *G. oxydans* than in exponential growth phase ^16^, suggesting LDT_Go_ expression might be upregulated when this bacterium transitions into growth arrest. However, this is not the case as LDT_Go_ expression and protein levels are comparable across growth phases (Fig. S17A).

Inspired by the above-mentioned inverse correlation between M4 and LD1,3-crosslinked dimers, we considered that the generation of 1,3-crosslinks could be modulated by monomer substrate availability. Enzymes controlling M4 abundance include DD-carboxypeptidases (DD-CPases) that cleave off the terminal D-Ala^5^, or DD-endopeptidases (DD-EPases), which break DD-crosslinked muropeptides (e.g., D44 dimers, crosslinked between D-Ala^4^-mDAP^3^) into M4 monomers (Fig 7A). As *G. oxydans* only encodes one putative DD-CPase (*gox0019)* and one DD-EPase with homology to PBP7 (*gox0607*), we generated individual deletion mutants and evaluated their implication on LD1,3-crosslink formation. Deletion of *gox0019* was not possible, suggesting this activity is essential in *G. oxydans*. On the contrary, the Δ*gox0607* strain was viable and exhibited a notable accumulation of DD-crosslinked peptides along with a decrease in M4 levels (Fig 7B, S17B). These findings align with its presumed function as a DD-EPase, and so we named it PBP7_Go_. Remarkably, Δ*pbp7_Go_* presented a ca. 50% reduction in the LD1,3-crosslink levels in stationary phase, effect that was reverted by genetic complementation (Fig 7C, S17B), confirming the implication of this enzyme in boosting the levels of LD1,3-crosslinks in stationary phase.

**Figure 7.**
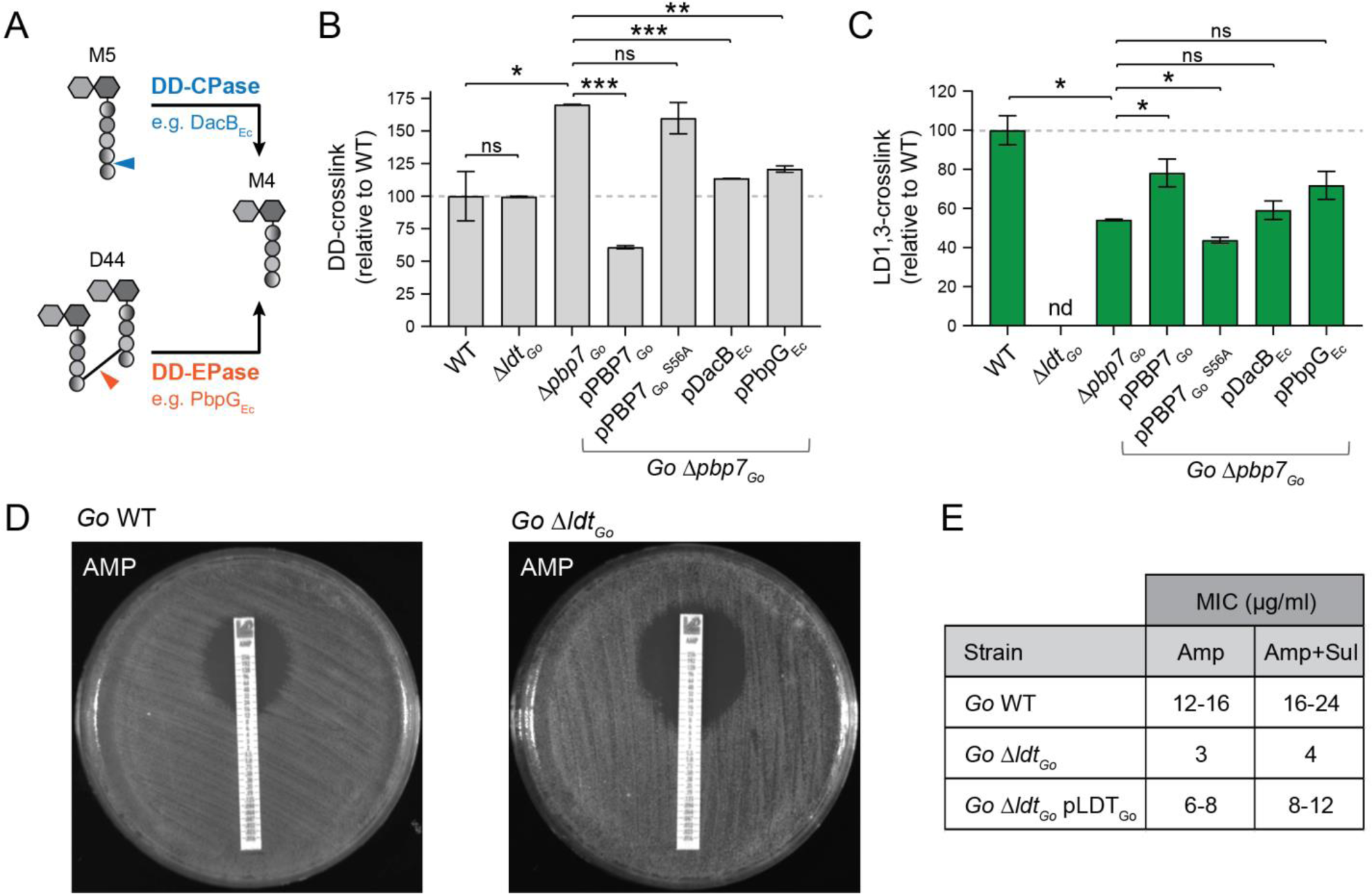
DD-crosslinking turnover controls LD1,3-crosslinking levels. A) Scheme of the production of M4 by DD-CPases (blue) or DD-endopeptidases (orange). Cleavage sites are indicated by arrowheads. DD-crosslinking (B) and LD-crosslinking (C) levels for *G. oxydans* (*Go*) WT, Δ*ldt_Go_*, Δ*pbp7_Go_* and the indicated complemented strains. Relative molar abundances of the DD- and LD1,3-crosslinked muropeptides were quantified and represented relative to the *Go* WT levels. D) Representative antibiograms of *G. oxydans* WT and Δ*ldt_Go_* with Ampicillin strips. F) Table indicating MIC values for Ampicillin (Amp) and Ampicillin/Sulbactam (Amp+Sul) for the *G. oxydans* WT, Δ*ldt_Go_* and Δ*ldt_Go_* pLDT_Go_ complemented strain. * p-value < 0.05; ** p-value < 0.001; *** p-value < 0.0001; ns: not significant.

To evaluate if the reduced LD1,3-crosslink levels in the Δ*pbp7_Go_* mutant is really the result of a reduction of the substrate flux or rather an effect on LDT_Go_ activity (e.g., the need of PBP7_Go_-LDT_Go_ protein-protein interactions), we generated a point mutant (S56A) in the predicted catalytic serine of PBP7_Go_. Expression of the allele carrying this point mutation did not complement the lower LD1,3-crosslinking of the Δ*pbp7_Go_*strain (Fig 7BC, S17B). Furthermore, the Δ*ldt_Go_* mutant strain presents DD-crosslinking levels that are comparable to *G. oxydans* wild-type strain, buttressing the idea that PBP7_Go_ DD-EPase feeds M4 to LDT_Go_ for LD1,3-crosslinking synthesis (Fig 7B, S17B). The sequential but independent action of PBP7_Go_ and LDT_Go_ was further confirmed by cross complementation of the Δ*pbp7_Go_* strain with alternative DD-EPases such as *E. coli* PbpG_Ec_ and DacB_Ec_. Expression of these ortholog endopeptidases reduced ca. 30% DD-crosslinking and concomitantly increased the LD1,3-crosslinking levels (Fig. 7BC, S17B). In sum, these results demonstrate a regulatory link between DD-crosslinking and LD-crosslinking synthesis.

Depletion of LD1,3-crosslinking has no phenotypical consequences in *G. oxydans* morphology or growth under optimal culture conditions (Fig. 7DE, S18AB). However, *G. oxydans* Δ*ldt_Go_* is approx. 4 times more sensitive than the wild-type strain when challenged with cell wall or membrane-active antibiotic such as ampicillin, fosfomycin and deoxycholate (Fig. 7DE, S18C). Interestingly, these phenotypes were alleviated by genetic complementation with LDT_Go_ (Δ*ldt_Go_* pLDT_Go_) but not when the *ldt_Go_* allele was replaced by YcbB_Ec_ (*ldt_Go_*::*ycbB_Ec_*) (Fig. S18C). In sum, these results suggest that LD1,3-crosslinks strengthen *G. oxydans* cell wall to withstand cell envelope perturbations and that they are not functionally exchangeable by 3,3 type LD-crosslinks.

## Discussion

The bacterial cell wall has been the subject of decades of research. Although several bacterial model organisms have been extensively investigated, less is known about how the cell wall is built and remodeled in other species. We found that the peptidoglycan of acetic acid bacteria was characterized for being amidated in the L-center of mDAP^3^ and for presenting a previously unrecognized LD-crosslink between the L-Ala^1^ and the DAP^3^ ^16^. Using *G. oxydans*, we report the identification, activity, and structural properties of this novel LD1,3-crosslinking enzymes.

Although LD1,3-TPases share a YkuD active domain with the well-studied LD3,3-TPases, these enzymes exhibit poor sequence identity and catalyze rather different crosslinks. Yet, our biochemical data suggested that these differences were likely not at the acceptor site given that, like YcbB_Ec_, LDT_Go_ can use muropeptides of varying peptide lengths such as tripeptides, tetrapeptides (e.g., Fig. 1A), and even pentapeptides (Fig. S19). Instead, structural differences in their donor site should explain how LD1,3-TPases make L-Ala^1^-DAP^3^ crosslinks and the unique nature of these enzymes in engaging non-terminal peptides in the crosslinking process, in stark contrast to the rigid specificity of LD3,3-TPases and DD-TPases, which exclusively operate on tetra and pentapeptide substrates, respectively.

Indeed, the donor site of the catalytic domain is very different between 1,3 and 3,3 LD-TPases. We identified a new region in our LDT_Go_ structure that consists of 2 interconnected loops between β-strands, which are broadly conserved among LD1,3-TPases. The structural properties of these loops shed light onto two important mechanistic questions: i) How does the acyl[enzyme complex occur? and, ii) how can these enzymes use different donor muropeptide substrates? The answer to the first question hides in one of the long linker loops. In LDT_Go_ and other Acetobacteria, this loop includes a trio of consecutive tyrosines that likely stabilize the glycanic moiety of the donor muropeptide, thereby bringing the L-Ala in position 1 of the peptide at just 3.3 Å from the catalytic Cys. In other LD1,3-TPases, this triad is often replaced by a DHF sequence; however, these residues could form the same glycan stacking interactions (Fig S15AB). Interestingly, networks of aromatic residues have been previously implicated in glycan stabilization in some endolysins of phages suggesting this strategy might also be used in other cell wall acting enzymes ^30^.

The interaction of the catalytic Cys with the L-Ala^1^ raises a dilemma: how do LD1,3-TPases accept longer muropeptides such as M4? In *Mycobacterium tuberculosis*, the Trp340 residue of the LD3,3-TPase LdtMt_2_ has been suggested to cause steric hindrances that restrict the enzyme’s substrate preference for tetrapeptides ^31^. Instead, LDT_Go_ has a large acidic cavity observed near the donor site that is compatible with the accommodation of muropeptides donor substrates of variable peptide length. This acidic character would prevent stabilization of the carboxylate-containing peptides at positions 2-4 (D-Glu^2^-mDAP^3^-D-Ala^4^) close to the catalytic Cys264 thus favoring interaction with L-Ala^1^.

The flexibility of LD1,3-TPases to accept various donor muropeptides might contribute to crosslinking homeostasis. It is known that several species increase their LD3,3-crosslinks in stationary phase ^9,32^. Similarly, LD1,3-crosslinking increases in *G. oxydans* and other Acetobacteria when the cultures reach high population densities ^16^. However, while such an upshift is often the result of a stress-driven upregulation of the enzyme ^9,31,33,34^, we found that in *G. oxydans,* LDT_Go_ levels are comparable across growth phases. Our results show instead that the endopeptidase PBP7_Go_ plays a critical role in regulating substrate availability for LD1,3-crosslinking (Fig. 7). Linking DD-crosslink turnover with LD1,3-crosslink synthesis could help *G. oxydans* peptidoglycan homeostasis, particularly during stationary phase of growth, when DD-transpeptidation declines due to reduced *de novo* synthesis and pentapeptide levels ^35^. Interestingly, the expression of PBP7_Go_ is not upregulated in stationary phase (Fig. S17C) suggesting a more complex regulation of the synthesis of LD1,3-crosslinks. To this end, we previously reported that the peptidoglycan of acetic acid bacteria is highly amidated in exponential phase, a modification that dramatically vanishes coinciding with the production of LD1,3-crosslinks in stationary phase ^16^. As this modification can negatively affect the activity of distinct endopeptidases ^16^, it is possible that in addition to the role of DD-endopeptidases as substrate suppliers, mDAP amidation might play a role in controlling the use of these monomers in LD1,3-crosslinking. Identification of the genetic determinants of L-mDAP amidation and its growth phase dependent regulation will likely provide new insights into crosslinking homeostasis of Acetobacteria.

An additional layer of complexity in the regulation of LD1,3-crosslinks emerges from the presence of a Pro-rich belt blocking the active site in our LDT_Go_ structure. This property, which was not predicted by the AF model, suggests a potential self-modulation of LDT_Go_ activity. Our results indicate that this protein is associated to the membrane *in vivo* and thus, it is unlikely that the belt needs to be cleaved to activate the protein. Instead, we hypothesize that a reversible binding of the belt can restrict or facilitate access of LDT_Go_ to the peptidoglycan layer in addition to unlocking its active site (Fig. S20). Along this line, it has been previously proposed that Pro-rich regions are associated with the bacterial cell wall ^36,37^ and N-terminal disordered extensions allow lipoproteins to cross the peptidoglycan and interact with their PBP partners ^38,39^. As the Pro-rich N-proximal belt does not seem to be restricted to Acetobacterial LDT_Go_ orthologs, we hypothesize that the reversible belt-inhibition of LD1,3-TPases should respond to general stimuli like those found in stationary phase (when LD1,3-crosslinking peaks up) rather than a specific lifestyle or environmental context.

Although LD1,3-crosslinks are not essential for *G. oxydans*, they do strengthen the cell wall integrity against diverse cell envelope aggression, including β-lactam antibiotics and membrane perturbations. Further, we previously demonstrated that this LD1,3-crosslink is immune to potential attacks caused by peptidoglycan-degrading predatory endopeptidases such as those delivered by type VI secretion systems ^16^. Based on our data, we propose a hypothesis that parallels the endogenous function of PBP7_Go_ as substrate supplier. We suggest that when exogenous DD-endopeptidases attack (or when PBPs are rendered inactive by β-lactams), the local availability of muropeptide monomers increases. Consequently, this would stimulate LD1,3-crosslinking activity, effectively repairing the damage by replacing the broken bond with an alternative, more resilient one within the peptidoglycan structure. As bacteria encoding LD1,3-TPases include opportunistic pathogens such as *Granulibacter bethesdensis* and *B. cenocepacia* ^40,41^, the activity of these enzymes might be a chemotherapeutic target to sensitize bacteria to β-lactams. Although it remains to be investigated whether the phenotypes observed for the *ldt_Go_* mutant are caused solely by a reduction of the peptidoglycan crosslinking levels or if potential interactions between the peptidoglycan and the outer membrane are also compromised, the observed the D,L-amino acid exchange activity of these enzymes supports this possibility. Future research will determine whether LD1,3-TPases can, like their Alphaproteobacteria LD3,3-TPase counterparts, tether the peptidoglycan to outer membrane proteins ^10,42^.

## Materials and methods

### Microbiology and Molecular Biology

All *Gluconobacter oxydans* strains used in this study were derived from the sequenced DSM-7145 ^43^. Strains used in this study are listed in Table S2. *G. oxydans* cultures were routinely grown aerobically to stationary phase in YP medium (5 gr Yeast extract, 5 gr peptone per liter) without or with 3% mannitol (YPM) at 30 ^°^C. *E. coli, V. cholerae* and *Burkholderia* strains were grown aerobically in Luria broth (LB) at 37 ^°^C. For agar plates, 15 g/l agar was added to the medium. Ampicillin (Amp) 100 µg/ml, Kanamycin (Kan) 50 µg/ml and Cefoxitine (Cfx) 50 µg/ml were used when required. In the amino acid exchange experiments, 10 mM D- or L-aa were used as supplements. Plasmids (Table S3) were constructed by standard DNA cloning techniques. Constructs were PCR-amplified and cloned into the plasmid pSEVA238, pET28b(+), pET22b(+) and pKOS6b plasmids as indicated in the text. Mutants in *G*. *oxydans* were constructed as described in ^44^ with some modifications. pKOS6b derivative plasmids carrying the 1Kbp flanking regions of the gene of interest were introduced by conjugation. Stationary phase *G. oxydans* recipient strain and S17-1 λ-PIR donor strains were washed of antibiotics, mixed in equal ratios, and placed on a YPM agar plate in a drop. After 24 h incubation, the cells were washed from the plate with fresh YPM medium and the resulting cell suspension was plated on YPM plates containing Kanamycin 50 µg/ml and Cefoxitine 50 µg/ml, as *G. oxydans* possesses a natural resistance to this antibiotic. Ectopic complementation of *G. oxydans* mutant strains was performed using pSEVA238 derivatives listed in Table S3, transferred by conjugation from S17-1 λ-PIR donor strains as described above. Induction of the gene expression was achieved by addition of benzoate 1 mM. For peptidoglycan analysis of the LDT activities *in vivo* in *E. coli*, BL21 (DE3) cultures transformed with the corresponding pET22b(+) and pET28b(+) plasmid derivatives (Tables S2 and S3) were grown to OD_600_ 0.4 units and induced with 1 mM IPTG during 3h.

### Growth curves, viability assays and antibiograms

For growth curves, stationary phase cultures were normalized to an OD_600_ of 0.04 and 20 μl were used for inoculating 96-well plates containing 180 μl of fresh YPM medium. At least three replicates per strain and condition were inoculated in two independent experiments. Optical density was monitored using an Eon Biotek plate reader (Biotek, Winooski, VT, USA), at 5 min intervals at 30 ^°^C. Viability assays were done with normalized overnight cultures subjected to serial 10-fold dilution. Five-microliter drops of the dilutions were spotted onto the indicated agar plates and incubated at the appropriate temperature for 24-48 h prior to image acquisition. For antibiograms, O.N cultures of the different strains were normalized to OD_600_: 0.05 and spread with a sterile cotton swap over a YPM agar plate. After the plates were air-dried, the MIC Test Strips (Liofilchem, Italy) were added. Plates were incubated for 24-48h at 30 °C and later inhibitory halo was quantified following manufacturing instructions.

### Protein expression and purification

The *G. oxydans, B. cenocepacia* and *E. coli* genes encoding LDT_Go_, LDT_Go_ C264A, LDT_Bcn_, LDT_Bcn_ C354A, LdcA_Ec_ ^24^, LdtA_Vc_ ^9^ were cloned on pET28b(+) (Novagen) with C-terminal His-tags for expression in *E. coli* BL21(DE3) cells ^45^. Bacteria were cultured in Terrific Broth (24 g/l Yeast extract, 20 g/l tryptone, 4 ml/l glycerol, 0.017 M KH_2_PO_4_, 0.072 M K_2_HPO_4_) and expression was induced at OD_600_=0.4 with 1 mM IPTG and left overnight at 16 ^°^C. Cell pellets were resuspended in PBS with a Complete Protease Inhibitor Cocktail Tablet (Roche) and lysed by 2 passes through a French press at 10,000 psi. After centrifugation (30 min, 100,000 *x g*), LdcA_Ec_ was purified from the cleared lysates via Ni-NTA agarose columns (Qiagen) and eluted with a discontinuous imidazole gradient using an ÄktaGo system. For LDT_Go_, LDT_Go_ C264A, and LDT_Bcn_, the pellet was treated with 0.1% (v/v) Triton X-100 overnight. The solubilized fraction was centrifuged (30 min, 100,000 *x g*) and the supernatant was purified via Ni-NTA agarose columns (Qiagen) and eluted with a discontinuous imidazole gradient using an ÄktaGo system, with buffers containing 0.1% (v/v) Triton X-100. For purification of PelB-LDT_Go_, used for crystallographic studies, a C-terminal 6His-tagged version was cloned onto pET22b (Novagen) for expression in *E. coli* BL21(DE3) cells using the signal peptide PelB sequence encoded in the plasmid. Cell pellets were resuspended in sodium phosphate buffer (PBS) with a Complete Protease Inhibitor Cocktail Tablet (Roche) and lysed using French press at 10,000 psi. After centrifugation (30 min, 100,000 *x g*, the cleared lysate was purified via Ni-NTA agarose columns (Qiagen) and eluted with a discontinuous imidazole gradient using an ÄktaGo system. Purified fractions were loaded on a size exclusion chromatography (SEC) Superdex 200 Increase 10/300 GL column equilibrated in 100 mM citrate/citric acid buffer pH 5 with 300 mM NaCl. Purified proteins were visualized by SDS-PAGE electrophoretic protein separation and quantified by Bio-Rad Protein Assay (Bio-Rad). The proteins were either stored at 4 °C for immediate use, or at −80 °C after addition of 10% (v/v) glycerol.

### Peptidoglycan isolation

Cells from 0.2 l cultures of overnight stationary phase or 1 L exponential phase (OD_600_ = 4.0) were pelleted at 5,250 *x g* and resuspended in 5 ml of PBS, added to an equal volume of 10% SDS in a boiling water bath and vigorously stirred for 4 h, then stirred overnight at room temperature. The insoluble fraction (peptidoglycan) was pelleted at 400,000 *x g*, 15 min, 30 °C (TLA-100.3 rotor; OptimaTM Max ultracentrifuge, Beckman) and resuspended in Milli-Q water. This step was repeated 4-5 times until the SDS was washed out. Next, peptidoglycan was treated with Pronase E 0.1 mg/ml at 60 °C for 1 h and then boiled in 1% SDS for 2 h to stop the reaction. After SDS was removed as described previously, peptidoglycan samples were resuspended in 200 μL of 50 mM sodium phosphate buffer pH 4.9 and digested overnight with 30 μg/ml muramidase (from *Streptomyces albus*) at 37 ^°^C. Muramidase digestion was stopped by heat-inactivation (boiling during 5 min). Coagulated protein was removed by centrifugation (20,000 *x g*, 15 min). The supernatants (soluble muropeptides) were subjected to sample reduction. First, pH was adjusted to 8.5-9 by addition of borate buffer 0.5M pH 9 and then muramic acid residues were reduced to muramitol by sodium borohydride treatment (NaBH_4_ 10 mg/ml final concentration) during 30 min at room temperature. Finally, pH was adjusted to 2.0-4.0 with orthophosphoric acid 25% prior to analysis by LC.

### Peptidoglycan analysis

Chromatographic analyses of muropeptides were performed by Ultra Performance Liquid Chromatography (UPLC) on an UPLC system (Waters) equipped with a trapping cartridge precolumn (SecurityGuard ULTRA Cartridge UHPLC C18 2.1 mm, Phenomenex) and an analytical column (BEH C18 column (130 Å, 1.7 μm, 2.1 mm by 150 mm; Waters, USA). Muropeptides were detected by measuring the absorbance at 204 nm using an ACQUITY UPLC UV−visible Detector. Muropeptides were separated using a linear gradient from buffer A (Water + 0.1 % (v/v) formic acid) to buffer B (Acetonitrile 100 % (v/v) + 0.1 % (v/v) formic acid) over 15 min with a flowrate of 0.25 ml/min. Individual muropeptides were quantified from their integrated areas using samples of known concentration as standards. Identity of the muropeptides was confirmed by MS-MS/MS analysis, using a Xevo G2-XS Q-tof system (Waters Corporation, USA). The instrument was operated in positive ionization mode. Detection of muropeptides was performed by MS^E^ to allow for the acquisition of precursor and product ion data simultaneously, using the following parameters: capillary voltage at 3.0 kV, source temperature to 120 ^°^C, desolvation temperature to 350 ^°^C, sample cone voltage to 40 V, cone gas flow 100 l/h, desolvation gas flow 500 l/h and collision energy (CE): low CE: 6 eV and high CE ramp: 15-40 eV. Mass spectra were acquired at a speed of 0.25 s/scan. The scan was in a range of 100–2000 m/z. Data acquisition and processing was performed using UNIFI software package (Waters Corp.). The quantification of muropeptides was based on their relative abundances (relative area of the corresponding peak) and relative molar abundances, as indicated elsewhere ^46^. A table of all the identified muropeptides and the observed ions is provided (Table S5).

### *In vitro* activity assays

The LDT_Go_, LDT_Bcn_, LDT_Bcn_ C354A, LdtA_Vc_, LdcA_Ec_ and KP27 reactions were performed in 50 µl reactions using 0.1 mg/ml of purified enzymes with 1 mg/ml of sacculi isolated from stationary phase cultures from *G. oxydans, G. oxydans* Δ*ldt_Go_*, *V. cholerae* WT or Δ*dacA1* mutant. To generate M3 and D-Met labeled sacculi, LdcA_Ec_ or LdtA_Vc_ with 10 mM of D-methionine (D-Met), respectively, were incubated for 2h at 37 °C prior to heat-inactivation and addition of LDT_Bcn_. LD1,3-TPase reactions were carried out in LD buffer (50 mM Tris HCl, pH 8, 50 mM NaCl) overnight at 30 ^°^C. For control reactions with KP27, KP buffer was used (20 mM Tris-HCl, pH 8, 1 mM MgCl_2_, 1 mM ZnCl_2_) at 37 °C for 90 minutes. Reactions were heat-inactivated, and fractions were separated by centrifugation at 20,000 *x g* for 15 min. For analysis of the insoluble product (pellet), reactions were finally treated with 50 µg/ml muramidase for 2h at 37 ^°^C, heat-inactivated and, followed by centrifugation at 20,000 *x g* for 15 min to remove precipitated proteins. Samples were subjected to reduction, pH adjusted, and analyzed by LC as described above. Analysis of the soluble fraction of the LD1,3-TPase in vitro reactions was performed for identification of released tripeptides. Non-reduced samples were pH adjusted and analyzed by LC-MS, as described above. To determine LDT_Bcn_ inhibition, Ampicillin (100 µg/ml), Imipenem (100 µg/ml) and Copper (1 mM) were added to the reaction before the addition of the enzyme.

#### Crystallization and structure determination

Purified LDT_Go_ was concentrated on a 10 kDa cutoff Amicon Ultra Centrifugal Filter (Merck-Millipore) to ∼10 mg/ml and loaded on a Superdex-200 increase 10/300 GL column equilibrated in 100 mM citrate/citric acid buffer pH 5.0, 300 mM NaCl. Protein peak fractions were concentrated further to 19 mg/ml. Crystals were grown at 20 °C by sitting drop vapor diffusion, using a 1:1 protein to reservoir ratio, in the A1 condition from the Morpheus screen, which contains 30 mM Magnesium chloride hexahydrate; 30 mM Calcium chloride dihydrate, 0.1M Imidazole; MES monohydrate (acid) pH 6.5 and 20% v/v PEG 500* MME; 10 % w/v PEG 20000. Crystals first appeared after ∼3 days, were fished after 3 weeks and flash-frozen in liquid nitrogen. X-ray diffraction data was collected on the ID23-2 beamline at the European Synchrotron Radiation Facility, Grenoble, France ^47^. The data was processed using XDS ^48^. The crystal belonged to P2_1_2_1_2_1_ space group and contained a single molecule in the asymmetric unit. The crystallographic phase problem was solved by using the AlphaFold2 model, generated in Colabfold ^49^ as search model in molecular replacement. Coot ^50^ as used to build the model and the structure was refined using Refmac5^51^ and PHENIX refine ^52^. For complete data collection and refinement statistics, see Table 1. The final model was validated using MolProbity ^53^. Atomic coordinates and structure factors of the LDT_Go_ structure have been deposited in the Protein Data Bank (PDB ID: 8QZG).

**Table 1.** Data collection and refinement statistics.

|  | **LDT<sub>Go</sub>** |
| --- | --- |
| Wavelength | 0.8731 |
| Resolution range | 48.41 - 1.732 (1.794 - 1.732) |
| Space group | P 21 21 21 |
| Unit cell | 52.778 Å, 56.688 Å, 93.019 Å<br>90°, 90°, 90° |
| Total reflections | 163948 (7995) |
| Unique reflections | 54354 (4155) |
| Multiplicity | 3.0 (1.9) |
| Completeness (%) | 98.48 (87.39) |
| Mean I/sigma(I) | 6.68 (0.74) |
| Wilson B-factor | 24.55 |
| R-merge | 0.1004 (0.9055) |
| R-meas | 0.1206 (1.16) |
| R-pim | 0.06601 (0.7139) |
| CC1/2 | 0.995 (0.355) |
| CC* | 0.999 (0.724) |
| Reflections used in refinement | 29296 (2335) |
| Reflections used for R-free | 1530 (119) |
| R-work | 0.1706 (0.3252) |
| R-free | 0.2051 (0.3319) |
| CC1/2 | 0.995 (0.355) |
| CC* | 0.999 (0.724) |
| Number of non-hydrogen atoms | 2191 |
| macromolecules | 2056 |
| ligands | 2 |
| solvent | 133 |
| Protein residues | 263 |
| RMS (bonds) | 0.006 |
| RMS (angles) | 0.72 |
| Ramachandran favored (%) | 96.14 |
| Ramachandran allowed (%) | 3.86 |
| Ramachandran outliers (%) | 0.00 |
| Rotamer outliers (%) | 0.00 |
| Clashscore | 0.98 |
| Average B-factor | 27.10 |
| macromolecules | 26.71 |
| ligands | 34.35 |
| solvent | 32.91 |

Statistics for the highest-resolution shell are shown in parentheses.

#### Molecular dynamics (MD) simulations

Both the full-length AlphaFold model and the gap-filled X-ray crystal structure of LDT_Go_ were immersed in a rectangular box of TIP3P waters, neutralized with counterions, energy minimized, and subjected to MD simulation under periodic boundary conditions for up to 400 ns using the *pmemd.cuda* module of AMBER 22 (https://ambermd.org/), following a previously described protocol ^54^. The peptidoglycan strand was modeled using the structure reported by ^55^ as a template2 and manually docked into LDT_Go_ to provide a complex that was simulated under identical conditions. AMBER ff19SB and GLYCAM_06j parameters were assigned to peptide and glycan atoms, respectively. Electrostatic interactions were represented using the smooth particle mesh Ewald method ^56^ with a grid spacing of 1 Å and the cutoff distance for the non-bonded interactions was 9 Å.

#### Detection of LDT_Go_ by Western-Blot

For localization of LDT_Go_ in the different cellular compartments, *G. oxydans ldt_Go_::ldt_Go_6his* was grown in YPM medium to stationary phase (OD_600_ 3.0 units). The samples were lysed using a French press at 10,000 psi. The membrane and soluble fractions were separated by ultracentrifugation at 75,000 *x g* for 60 min. The membrane fractions were resuspended in 200 µl PBS and the soluble fraction was concentrated to a similar level using Amicon^R^ Ultra 15 centricons (Millipore). 15 µg of membrane and soluble fraction were loaded after normalization by total protein quantification using the Bio-Rad Protein Assay (Bio-Rad) onto a 12% acrylamide gel. For the quantification of LDT_Go_-His in the different growth phases, cultures of *G. oxydans ldt_Go_::ldt_Go_6his* were collected at OD_600_ 0.3 (Exp), 1.2 (Early Sta) and 3 (Late Sta). The samples were OD normalized and prepared as described above. For quantification, 15 µg of the membrane fraction was loaded onto the 12% acrylamide gel.

#### Beta-galactosidase assay

The presumed promoter region corresponding to the 500 bp upstream from the putative *gox1073-1074* operon, was cloned into the promoter probe plasmid pSEVA235, functional in *G. oxydans*. Beta-galactosidase activity of the promoter-lacZ translational fusion was measured through *o*-nitrophenyl-β-d-galactopyranoside (ONPG) cleavage by the product of the *lacZ* reporter gene, and specific activity was calculated in Miller units ^57^. Cultures of *G. oxydans* wild-type WT carrying pSEVA235-Pldt_Go_ derivative (and pSEVA235 empty plasmid as negative control) were grown in YPM medium at 30 °C to OD_600_ 0.3 units (Exp), 1.2 units (Early Sta) or 3 units (Late Sta) phase. Aliquots (100 μL) of three different subcultures were collected, and cells were permeabilized and assayed in triplicate for each strain as previously described ^57^.

#### Quantitative real-time PCR (RT-PCR)

The expression levels of *ldt_Go_* and *pbpb7_Go_*genes in exponential and stationary phase was analyzed by RT-PCR. RNA was isolated from *G. oxydans* wild-type cultures at OD_600_ 0.3 units (Exp) or 3 units (Sta) using the RNeasy Mini Kit (Qiagen), following the manufacturer’s instructions. Reverse transcription was performed with the High-Capacity cDNA Reverse Transcription Kit (Thermo Fisher), using aliquots of 5 µg total RNA. The synthesized cDNA was purified using a QIAquick PCR purification kit (Qiagen), and its concentration was determined spectrophotometrically in a Nano-drop Lite spectrophotometer (Thermo Scientific). Real time quantitative PCR (qRT-PCR) was performed using an iQ^TM^5 Multicolor Real-Time PCR Detection System (BIO-RAD) with qPCRBIO SyGreen Mix fluorescein (PCR BIOSYSTEMS). Master mixes were prepared as recommended by the manufacturer, with qRT-PCR primers listed in TableS4. Two independent experiments were carried out in triplicates for each data point. The relative quantification in gene expression was determined using the 2^-ΔΔCt^ method ^58^, using *recA_Go_ (gox1522)* gene control.

### Microscopy

For imaging, bacteria were immobilized on YPM pads containing 1% agarose. Phase-contrast microscopy was performed using a Zeiss Axio Imager.Z2 microscope (Zeiss, Oberkochen, Germany) equipped with a Plan-Apochromat × 63 phase-contrast objective lens and an ORCA-Flash 4.0 LT digital CMOS camera (Hamamatsu Photonics, Shizuoka, Japan), using the Zeiss Zen Blue software. Image analysis and processing were performed using Fiji ^59^ and the MicrobeJ plugins ^60^. For incorporation of FDAA, 1 mM of HADA ^25^ was added to each culture at exponential phase and incubated for 30 min. Cultures were then quenched with cold methanol in ice for 20 min and visualized as described.

#### Bioinformatic analyses

Orthologue sequences of LDT_Go_, corresponding to LD1,3-TPases, were searched by BLAST using the NCBI non-redundant protein database. Results were filtered using the following criteria: E-value > 10, coverage > 0.5, identity > 30. LD3,3-TPases (YcbB-like orthologues) were downloaded from OrthoDBv11 (LD-transpeptidase group 29809748at2) ^61^. For construction of a phylogenetic tree and conservation of proteins, we produced a tree of all species on OrthoDBv11 using PhyloT and orthologues of LD1,3- or LD3,3-TPases were mapped against this. The final tree was visualized using iTOL ^62^. Multisequence alignments were performed with Clustal Omega ^63^ or T-COFFE Expresso ^64^. Alignments were visualized and analyzed using Jalview v2 ^65^. We used ESPript for rendering sequence similarities and secondary structure information from aligned sequences, using the LDT_Go_ crystal structure (PDB ID: 8QZG) as reference ^66^. Signal peptide predictions were performed with SignalP 6.0^17^. Sequence logos were generated in R v4.3 using the ggseqlogo package ^67^.

### Statistical analysis

Statistical significance was assessed using the Student’s *t*-test. A p-value of less than 0.05 was considered statistically significant. Assays were performed with three biological replicates unless otherwise indicated.

## Acknowledgements

We thank the Cava lab members for insightful discussions. We also thank David Kostner for generously providing the plasmid pKOS6b. Research in the Cava lab was supported by the Swedish Research Council (2018-02823 and 2018-05882), Umeå University, the Knut and Alice Wallenberg Foundation and the Kempe Foundation (SMK2062). We acknowledge MAX IV Laboratory for time on Beamline BioMax under Proposal 20190808. Research conducted at MAX IV, a Swedish national user facility, is supported by the Swedish Research council under contract 2018-07152, the Swedish Governmental Agency for Innovation Systems under contract 2018-04969, and Formas under contract 2019-02496. We acknowledge the European Synchrotron Radiation Facility for provision of synchrotron radiation facilities (beamline ID23-2). Research in the Berntsson lab was supported by grants from the Swedish Research Council (2016-03599), the Knut and Alice Wallenberg Foundation, and the Kempe Foundation (SMK-1762 & SMK-1869). Research in the Gago lab is funded by the Spanish MICINN Projects PID2019-104070RB-C22 and AEI/10.13039/501100011033.

## Legends to Supplementary Figures

**Figure S1. Candidate YkuD-domain containing enzyme Gox2269**. A) Domain structure of the protein. B) UV muropeptide profile of *G. oxydans* wild-type (WT) and Δ*gox2269* mutant. LD1,3-crosslinked muropeptides are highlighted in green.

**Figure S2. LDT_Go_ signal peptide.** A) LDT_Go_ is predicted to have an SPII lipobox (Sec/SPII). B) Detection of C-terminal His-tagged LDT_Go_ in the particulate (Mb; membrane) or soluble fraction (S; cytoplasm and periplasm). Protein amount is normalized by total protein amount. C) Scheme of the LDT_Go_ WT and derivatives used in panel D: one lacking its signal peptide (ΔSP) and the second replacing it by YcbB_Ec_ signal peptide. Amino acid positions within LDT_Go_ protein sequence are indicated. D) LD1,3-crosslinking quantifications and E) UV muropeptide profiles of *G. oxydans* Δ*ldt_Go_*strain complemented with LDT_Go_ WT and derivatives indicated in panel C.

**Figure S3. LD1,3-crosslinking activity in *Burkholderia*.** A) UV muropeptide profiles of the indicated *Burkholderia* species. LD1,3-crosslinked dimers D13^NH^^2^ and D14 are highlighted in green. B, C) MS extracted ion current (XIC) traces of the indicated muropeptides.

**Figure S4. Detection of LD1,3-crosslinked muropeptides *in vitro.*** MS extracted ion current (XIC) trace of the D14 muropeptide in the *in vitro* activity assays of LDT_Bcn_ on M4-rich peptidoglycan sacculi (from *V. cholerae*), LDT_Bcn_ C354A point mutant (negative control) and assays with added Ampicillin 100 µg/ml (Amp), Imipenem 100 µg/ml (Imp) and copper 1 mM (Cu^2+^).

**Figure S5. Detection of tripeptide *in vitro***. A) Scheme of KP27 endopeptidase reaction on peptidoglycan sacculi. KP27 cleaving site between L-Ala^1^ and D-Glu^2^ is indicated with a blue arrowhead. B) UV muropeptide profiles of sacculi incubated (KP27) or not (control) with KP27 prior digestion with muramidase to release individual muropeptides. KP27(SN) corresponds to the analysis of free peptidoglycan soluble fragments released to the supernatant after KP27 digestion, which mostly include linear peptidoglycan chains of M1, and free tripeptides as illustrated in A. The MS extracted ion current (XIC) traces of the liberated tripeptide are shown.

**Figure S6. LDT_Bcn_ activity on peptidoglycan tripeptides.** A) Scheme of the preparation of M3-enriched peptidoglycan sacculi using the LD-CPase activity of LdcA of *E. coli*. B) UV muropeptide profiles, D13 MS extracted ion current (XIC) trace and C) quantification of the relevant muropeptides indicated. Variation is calculated as difference in relative molar abundance of the muropeptide in the LDT_Bcn_ vs control in the *in vitro* assays.

**Figure S7. Incorporation of D-Met into the peptidoglycan of *G. oxydans*.** UV muropeptide profiles from cultures of *G. oxydans* WT, Δ*ldt_Go_* and a derivative strain in which the *ldt_Go_* allele is replaced by *ycbB_Ec._* (*ldt_Go_::ycbB_Ec_*) supplemented or not (Control) with 10 mM of D-Met. LD1,3-crosslinked muropeptides are highlighted in green and the D-Met-modified M4 (M4^Met^) in blue.

**Figure S8. Identification of M2^X^ muropeptides in *G. oxydans* peptidoglycan.** Detection of the parental ion (left), its fragmentation pattern (middle) and determined structure based on the MS/MS data for A) M2^Glu^, B) M2^Gln^, C) M2^Phe^, and D) M2^Trp^.

**Figure S9. Functional association between M2^X^ muropeptides and LD1,3-TPases.** A) UV muropeptide profiles and the M2^X^ MS extracted ion current (XIC) traces of the indicated muropeptides in *G. oxydans* wild-type (WT), Δ*ldt_Go_* mutant and complemented strains. (B) UV muropeptide profiles and the indicated M2^X^ MS extracted ion current (XIC) traces of the indicated muropeptides in *E. coli* BL21 expressing LDT_Bcn_ or its catalytically inactive derivative C354A.

**Figure S10. Detection of M2^Phe^ *in vitro*.** UV muropeptide profiles and zoom-in of *in vitro* amino acid exchange reactions using *G. oxydans* WT peptidoglycan sacculi as substrate +/- 10 mM of D- or L-Phe, incubated with LDT_Bcn_. The M2^Phe^ MS extracted ion current (XIC) traces are indicated.

**Figure S11. *In vitro* activity of crystallized LDT_Go_.** Muropeptide profile of *E. coli* (*Ec*) BL21 expressing PelB-LDT_Go_ and extracted ion of the D14 muropeptide. The N-terminal PelB leader sequence from the pET22b(+) plasmid directs the protein to the periplasmic space.

**Figure S12. Conservation of the Pro-rich belt.** A) Domain analysis of representative LDT_Go_-like proteins from different bacterial taxa. Lipoprotein signal peptide SPII (Sec/SPII) is indicated in orange, translocation signal peptide (SPI/SPI) in blue, transmembrane domain ™ in purple, the Pro-rich belt in red, and the YkuD domain in grey. Prediction of the signal peptides was performed using SignalP 6.0. B) Multiple sequence alignment of the indicated LDT_Go_-like proteins from different bacterial species. Secondary structure of LDT_Go_ is shown immediately above the alignment. Beta sheets and alpha helices are indicated in blue and green, respectively, and the conserved Cysteine and Histidine catalytic residues are indicated with orange arrows. Colors in the sequence alignment are as described in A.

**Figure S13. Active state model of LDT_Go_.** A) Root-mean-square fluctuation (RMSF) during the 250 ns MD simulations of the crystallographic structure. Regions experiencing larger fluctuations are indicated in the orange boxed areas and labeled. B) Comparison between the LDT_Go_ AlphaFold2 (AF2, light brown) model and the molecular dynamics simulation (MDS, blue) model of LDT_Go_ lacking the belt. Relevant loops and catalytic residues are indicated.

**Figure S14. Distinctive active site organization between 3,3 and 1,3 LD-TPases.** A) Comparative overview between YcbB_Ec_ (blue) bound to meropenem (yellow) and the Alphafold2 model of LDT_Go_, (AF2, light brown). B) Zoom-in view of the active site. Relevant loops and catalytic residues are indicated.

**Figure S15. Sequence and structure identity between LDT_Go_-like proteins.** A) Conservation of the triad predicted to interact with NAM moiety at the donor site of LDT_Go_. Zoom-in view of the triad organization between the LDT_Go_ structure (green) which displays a YYY triad and the LDT_Bcn_ Alphafold2 model (AF2, blue) which displays a DHF triad. C) Comparative sequence analysis of the listed LDT_Go_ orthologs. Relevant conserved residues important substrate stabilization are indicated. D-I) Comparison of Alphafold2 models of the indicated LDT_Go_ orthologs. Zoom-in view of the donor site of the active domain is shown. Relevant loops and residues are indicated.

**Suppl Figure 16. Modelization of a large PG chain attached to the active site of LDT_Go_.** A) Molecular surface of LDT _Go_ (green) with the catalytic C264 highlighted in yellow. The PG layer is represented as capped sticks (C atoms colored in dark red). The positions of the relevant loops are indicated. B) Detailed view of the interaction of the PG chain in the LDT_Go_ active site. Relevant residues in the protein are represented as capped sticks and labeled. While the interactions are preserved for the NAM-peptide moiety close to the catalytic C264 residue (see Figure 6), expansion to a larger PG chain shows that the capping loop and L247-252 also play a relevant role in stabilization of the glycan chain far from the active site by the aromatic residues Y207 (from the capping loop) and the F251 (from L247-252).

**Figure S17. PBP7_Go_ controls the *in vivo* levels of LD1,3-crosslinking.** A) Detection of C-terminal His-tagged LDT_Go_ protein levels by Western blot and transcriptional levels by beta-galactosidase activity (Miller units) assays from cultures collected at exponential (Exp), early stationary (Early Sta) and stationary (Sta) growth phase as indicated in the methods section. Protein amount is normalized by total protein amount. B) UV muropeptide profiles of *G. oxydans* WT, Δ*ldt_Go_*, Δ*pbp7_Go_* and complemented strains as indicated. C) Quantitative RT-PCR analysis of *ldt_Go_*and *pbp7_Go_* genes in exponential (Exp) and stationary (Sta) growth phases. The house keeping gene *recA_Go_* is used as control. Data from three replicas in two independent experiments was analyzed using the 2^-ΔΔCt^ analysis method.

**Figure S18. Phenotypic characterization of the *G. oxydans*** Δ***ldt_Go_* mutant.** A) Phase contrast microscopy of wild-type (WT) and Δ*ldt_Go_* mutant. Scale bar: 2 µm. Cell length and mean width of the WT (n = 703) and Δ*ldt_Go_* mutant (n = 1587) are indicated. No significant differences found, pValue = 0.8864, applying non-parametric Mann-Whitney U test. B) Growth curves of the WT (grey) and Δ*ldt_Go_* mutant (green) grown in YPM and YP medium. B) Serial dilutions (10^−1^ to 10^−6^) from overnight cultures of wild-type (WT), Δ*ldt_Go_* mutant and allelic exchange with YcbB_Ec_ (*ldt_Go_::ycbB_Ec_*) were spotted onto YPM agar plates supplemented with Ampicillin 65 µg/ml (AMP), deoxycholate 0.0001 % (w/v) (DOC) and Fosfomycin 5 µg/ml (FOS). Growth on non-supplemented plate (YPM) was used as control.

**Figure S19. LDT_Bcn_ activity on peptidoglycan pentapeptides.** A) Scheme of the DacA1 DD-CPase activity and the muropeptide architecture of the peptidoglycan of *V. cholerae* WT and its Δ*dacA1* mutant derivative strain (M5-enriched). B) UV muropeptide profiles of the *V. cholerae* WT sacculi (M4 rich) and the Δ*dacA* mutant (M5-enriched, pentapeptide muropeptides highlighted in purple) treated or not (Control) with LDT_Bcn_. The LD1,3-crosslinked dimer D15 is highlighted in green, and its MS extracted ion current (XIC) trace is shown in the right-side panel.

**Figure S20. Model of the regulation of LDT_Go_ by the Pro-rich belt.** Graphic illustration suggesting how the detachment of the belt would facilitate both unblocking the active site and approaching it to the peptidoglycan.

## Supplementary tables

**Table S1.** Hydrogen bonds between the Pro-rich belt and the catalytic center.

| Pro-rich belt residue: | H-bond with residue: | Distance *: |
| --- | --- | --- |
| Pro56 (main-chain, =O) | Tyr181 (side-chain, OH) | 2.9 Å |
| Ala59 (main-chain, NH) | Glu262 (side-chain, =O) | 2.8 Å |
| Ser60 (main-chain, =O) | His245 (side-chain, NH) ** | 3.2 Å |
| Asn61 (side-chain, =O) | His245 (side-chain, NH) ** | 2.7 Å |
| Asn61 (side-chain, NH<sub>2</sub>) | Glu262 (side-chain, O<sup>-</sup>) | 3.1 Å |
| Thr64 (main-chain, =O) | Leu196 (main-chain, NH) | 3.0 Å |
| Thr64 (side-chain, OH) | Gly197 (main-chain, NH) | 2.8 Å |
| Thr66 (main-chain, NH) | Asp194 (main-chain, =O) | 2.9 Å |
| Thr66 (side-chain, OH) | Ala193 (main-chain, =O) | 2.9 Å |
| Pro67 (main-chain, =O) | Arg294 (side-chain, NH<sub>2</sub>) | 2.8 Å |
| Pro80 (main-chain, =O) | Gln287 (side-chain, NH<sub>2</sub>) | 2.8 Å |
| Pro83 (main-chain, =O) | Arg136 (side-chain, NH) | 2.7 Å |

\* As measured between the oxygen and the nitrogen/oxygen.
\*\* As both interactions are with the same nitrogen from the ring of the catalytic His245, only one of these will form at the time. Probably the interaction with Asn61 will dominate as it is over the shortest distance.

**Table S2.**
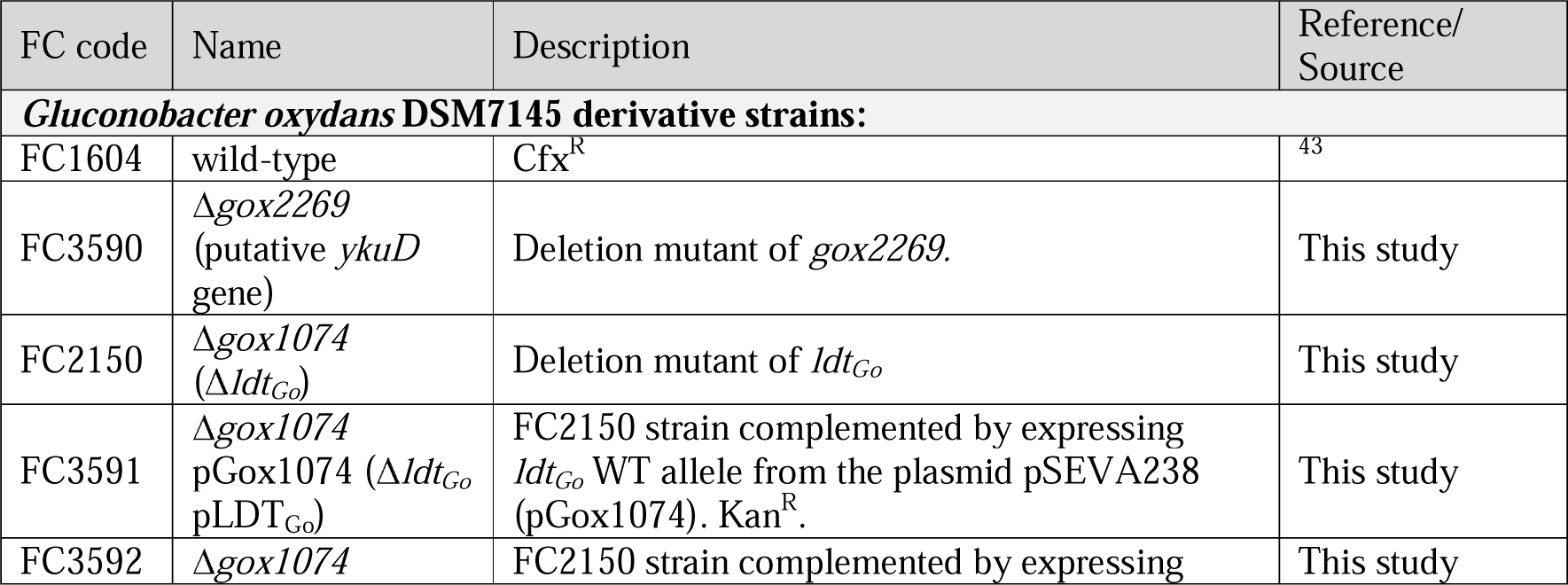
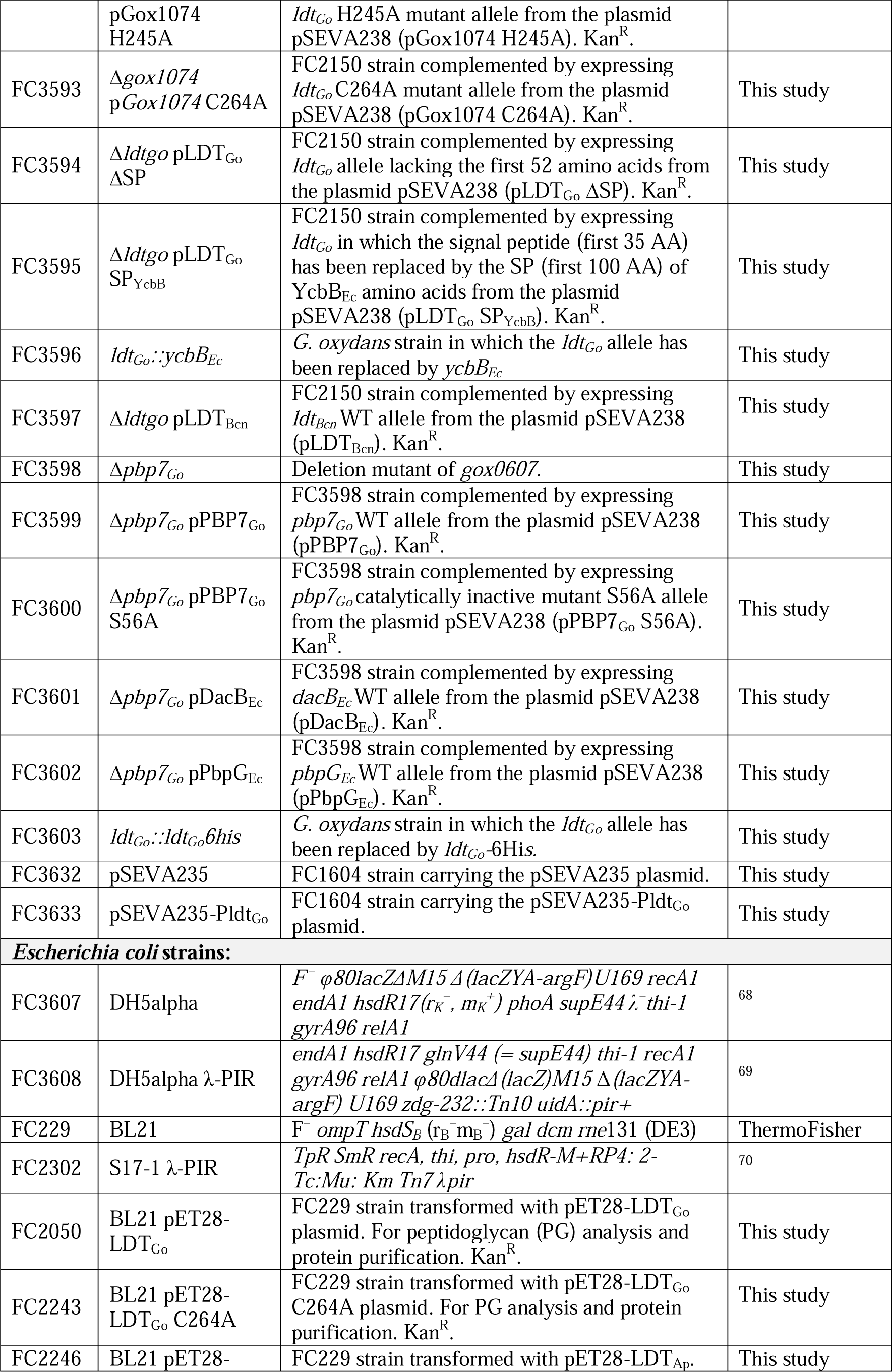
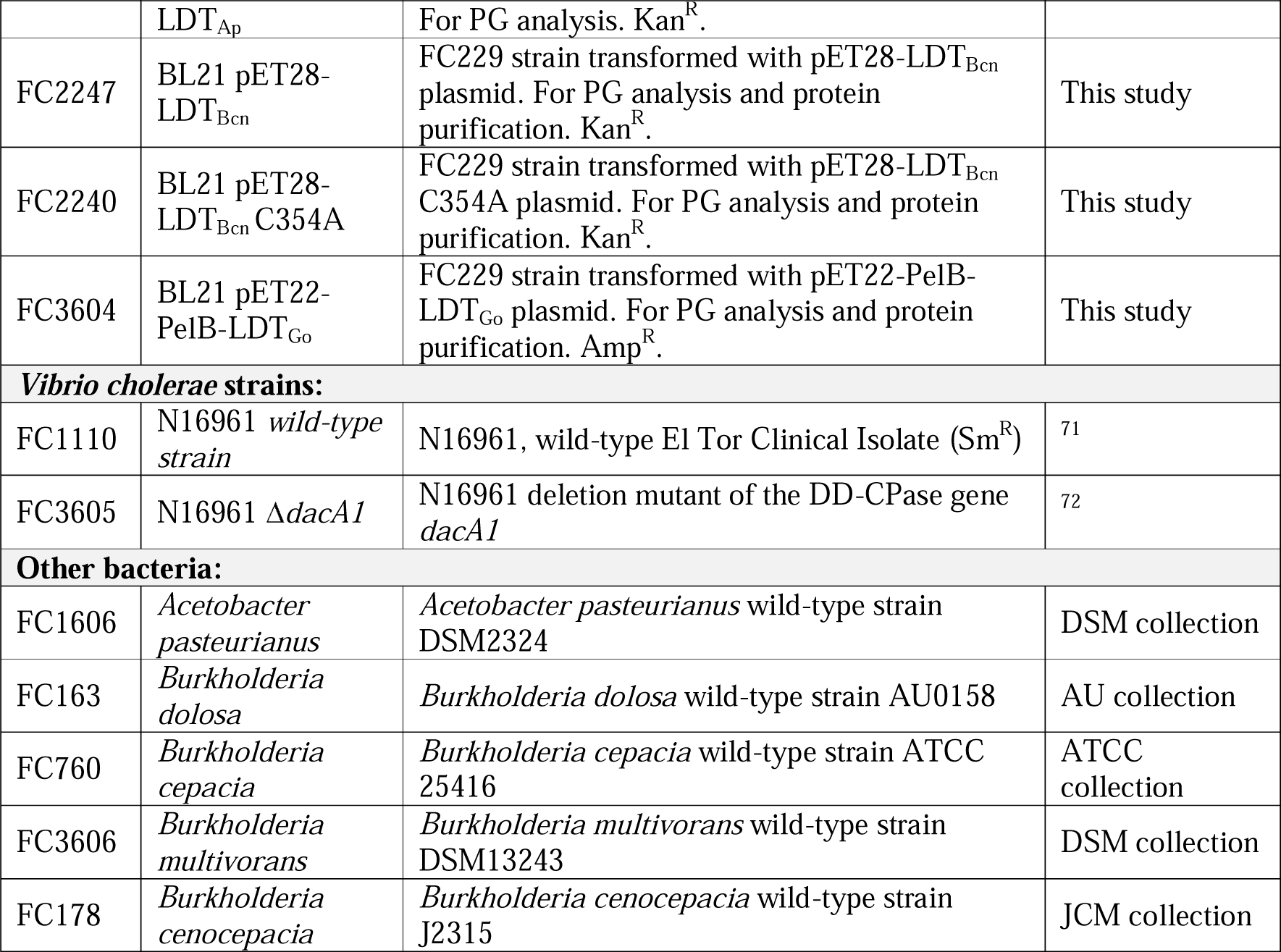
Strains.

| FC code | Name | Description | Reference/<br>Source |
| --- | --- | --- | --- |
| ***Gluconobacter oxydans* DSM7145 derivative strains:** |  |  |  |
| FC1604 | wild-type | Cfx<sup>R</sup> | 43 |
| FC3590 | Δ*gox2269*<br>(putative *ykuD*<br>gene) | Deletion mutant of *gox2269*. | This study |
| FC2150 | Δ*gox1074*<br>(Δ*ldt<sub>Go</sub>*) | Deletion mutant of *ldt<sub>Go</sub>* | This study |
| FC3591 | Δ*gox1074*<br>pGox1074 (Δ*ldt<sub>Go</sub>*<br>pLDT<sub>Go</sub>) | FC2150 strain complemented by expressing<br>*ldt<sub>Go</sub>* WT allele from the plasmid pSEVA238<br>(pGox1074). Kan<sup>R</sup>. | This study |
| FC3592 | Δ*gox1074* | FC2150 strain complemented by expressing | This study |
|  | pGox1074 H245A | *ldt<sub>Go</sub>* H245A mutant allele from the plasmid pSEVA238 (pGox1074 H245A). Kan<sup>R</sup>. |  |
| FC3593 | Δ*gox1074* pGox1074 C264A | FC2150 strain complemented by expressing *ldt<sub>Go</sub>* C264A mutant allele from the plasmid pSEVA238 (pGox1074 C264A). Kan<sup>R</sup>. | This study |
| FC3594 | Δ*ldtgo* pLDT<sub>Go</sub> ΔSP | FC2150 strain complemented by expressing *ldt<sub>Go</sub>* allele lacking the first 52 amino acids from the plasmid pSEVA238 (pLDT<sub>Go</sub> ΔSP). Kan<sup>R</sup>. | This study |
| FC3595 | Δ*ldtgo* pLDT<sub>Go</sub> SP<sub>YcbB</sub> | FC2150 strain complemented by expressing *ldt<sub>Go</sub>* in which the signal peptide (first 35 AA) has been replaced by the SP (first 100 AA) of YcbB<sub>Ec</sub> amino acids from the plasmid pSEVA238 (pLDT<sub>Go</sub> SP<sub>YcbB</sub>). Kan<sup>R</sup>. | This study |
| FC3596 | *ldt<sub>Go</sub>::ycbB<sub>Ec</sub>* | *G. oxydans* strain in which the *ldt<sub>Go</sub>* allele has been replaced by *ycbB<sub>Ec</sub>* | This study |
| FC3597 | Δ*ldtgo* pLDT<sub>Bcn</sub> | FC2150 strain complemented by expressing *ldt<sub>Bcn</sub>* WT allele from the plasmid pSEVA238 (pLDT<sub>Bcn</sub>). Kan<sup>R</sup>. | This study |
| FC3598 | Δ*pbp7<sub>Go</sub>* | Deletion mutant of *gox0607*. | This study |
| FC3599 | Δ*pbp7<sub>Go</sub>* pPBP7<sub>Go</sub> | FC3598 strain complemented by expressing *pbp7<sub>Go</sub>* WT allele from the plasmid pSEVA238 (pPBP7<sub>Go</sub>). Kan<sup>R</sup>. | This study |
| FC3600 | Δ*pbp7<sub>Go</sub>* pPBP7<sub>Go</sub> S56A | FC3598 strain complemented by expressing *pbp7<sub>Go</sub>* catalytically inactive mutant S56A allele from the plasmid pSEVA238 (pPBP7<sub>Go</sub> S56A). Kan<sup>R</sup>. | This study |
| FC3601 | Δ*pbp7<sub>Go</sub>* pDacB<sub>Ec</sub> | FC3598 strain complemented by expressing *dacB<sub>Ec</sub>* WT allele from the plasmid pSEVA238 (pDacB<sub>Ec</sub>). Kan<sup>R</sup>. | This study |
| FC3602 | Δ*pbp7<sub>Go</sub>* pPbpG<sub>Ec</sub> | FC3598 strain complemented by expressing *pbpG<sub>Ec</sub>* WT allele from the plasmid pSEVA238 (pPbpG<sub>Ec</sub>). Kan<sup>R</sup>. | This study |
| FC3603 | *ldt<sub>Go</sub>::ldt<sub>Go</sub>6his* | *G. oxydans* strain in which the *ldt<sub>Go</sub>* allele has been replaced by *ldt<sub>Go</sub>-6His*. | This study |
| FC3632 | pSEVA235 | FC1604 strain carrying the pSEVA235 plasmid. | This study |
| FC3633 | pSEVA235-Pldt<sub>Go</sub> | FC1604 strain carrying the pSEVA235-Pldt<sub>Go</sub> plasmid. | This study |
| **Escherichia coli strains:** |  |  |  |
| FC3607 | DH5alpha | *F<sup>-</sup> ϕ80lacZΔM15 $\Delta$(lacZYA-argF)U169 recA1 endA1 hsdR17(r<sub>K</sub><sup>-</sup>, m<sub>K</sub><sup>+</sup>) phoA supE44 λ-thi-1 gyrA96 relA1* | 68 |
| FC3608 | DH5alpha λ-PIR | *endA1 hsdR17 glnV44 (= supE44) thi-1 recA1 gyrA96 relA1 ϕ80dlacΔ(lacZ)M15 $\Delta$(lacZYA-argF) U169 zdg-232::Tn10 uidA::pir+* | 69 |
| FC229 | BL21 | *F<sup>-</sup> ompT hsdS<sub>B</sub> (r<sub>B</sub><sup>-</sup>m<sub>B</sub><sup>-</sup>) gal dcm rne131 (DE3)* | ThermoFisher |
| FC2302 | S17-1 λ-PIR | *TpR SmR recA, thi, pro, hsdR-M+RP4: 2-Tc:Mu: Km Tn7 λpir* | 70 |
| FC2050 | BL21 pET28-LDT<sub>Go</sub> | FC229 strain transformed with pET28-LDT<sub>Go</sub> plasmid. For peptidoglycan (PG) analysis and protein purification. Kan<sup>R</sup>. | This study |
| FC2243 | BL21 pET28-LDT<sub>Go</sub> C264A | FC229 strain transformed with pET28-LDT<sub>Go</sub> C264A plasmid. For PG analysis and protein purification. Kan<sup>R</sup>. | This study |
| FC2246 | BL21 pET28- | FC229 strain transformed with pET28-LDT<sub>Ap</sub>. | This study |
|  | LDT<sub>Ap</sub> | For PG analysis. Kan<sup>R</sup>. |  |
| FC2247 | BL21 pET28-LDT<sub>Bcn</sub> | FC229 strain transformed with pET28-LDT<sub>Bcn</sub> plasmid. For PG analysis and protein purification. Kan<sup>R</sup>. | This study |
| FC2240 | BL21 pET28-LDT<sub>Bcn</sub> C354A | FC229 strain transformed with pET28-LDT<sub>Bcn</sub> C354A plasmid. For PG analysis and protein purification. Kan<sup>R</sup>. | This study |
| FC3604 | BL21 pET22-PelB-LDT<sub>Go</sub> | FC229 strain transformed with pET22-PelB-LDT<sub>Go</sub> plasmid. For PG analysis and protein purification. Amp<sup>R</sup>. | This study |
| ***Vibrio cholerae* strains:** |  |  |  |
| FC1110 | N16961 *wild-type strain* | N16961, wild-type El Tor Clinical Isolate (Sm<sup>R</sup>) | 71 |
| FC3605 | N16961 Δ*dacA1* | N16961 deletion mutant of the DD-CPase gene *dacA1* | 72 |
| **Other bacteria:** |  |  |  |
| FC1606 | *Acetobacter pasteurianus* | *Acetobacter pasteurianus* wild-type strain DSM2324 | DSM collection |
| FC163 | *Burkholderia dolosa* | *Burkholderia dolosa* wild-type strain AU0158 | AU collection |
| FC760 | *Burkholderia cepacia* | *Burkholderia cepacia* wild-type strain ATCC 25416 | ATCC collection |
| FC3606 | *Burkholderia multivorans* | *Burkholderia multivorans* wild-type strain DSM13243 | DSM collection |
| FC178 | *Burkholderia cenocepacia* | *Burkholderia cenocepacia* wild-type strain J2315 | JCM collection |

**Table S3.** Plasmids.

| FC code | Name | Description | Reference |
| --- | --- | --- | --- |
| FC3610 | pKOS6b | Suicide vector for construction of deletion mutants in *G. oxydans*. Kan<sup>R</sup>. | 44 |
| FC3611 | pKOS6b-*gox2269* | pKOS6b derivative for clean deletion of *gox2269*. Kan<sup>R</sup>. | This study |
| FC2095 | pKOS6b-*gox1074* | pKOS6b derivative for clean deletion of *gox1074* (*ldt<sub>Go</sub>*). Kan<sup>R</sup>. | This study |
| FC3612 | pKOS6b-*ldt<sub>Go</sub>::ldt<sub>Go</sub>6his* | pKOS6b derivative for allelic exchange of the *ldt<sub>Go</sub>* locus with a 6-His tagged version. Kan<sup>R</sup>. | This study |
| FC3613 | pKOS6b-*gox0607* | pKOS6b derivative for clean deletion of *gox0607* (*pbp7<sub>Go</sub>*). Kan<sup>R</sup>. | This study |
| F3614 | pKOS6b-*ldt<sub>Go</sub>::ycbB<sub>Ec</sub>* | pKOS6b derivative for allelic exchange of the *ldt<sub>Go</sub>* locus with *ycbB* from *E. coli*. Kan<sup>R</sup>. | This study |
| FC2626 | pSEVA238 | Plasmid used for complementation in *G. oxydans*. Expression of genes under the control of the xylS-Pm promoter. Activated by benzoate or m-toluate. Kan<sup>R</sup> | 73 |
| FC3615 | pGox1074 (pLDT<sub>Go</sub>) | pSEVA238 derivative for expression of Gox1074 (LDT<sub>Go</sub>). Kan<sup>R</sup>. | This study |
| FC3616 | pGox1074 H245A | pSEVA238 derivative for expression of Gox1074 (LDT<sub>Go</sub>) H245A point mutant. Kan<sup>R</sup>. | This study |
| FC3617 | pGox1074 C264A | pSEVA238 derivative for expression of Gox1074 (LDT<sub>Go</sub>) C264A point mutant. Kan<sup>R</sup>. | This study |
| FC3618 | pLDT<sub>Go</sub> ΔSP | pSEVA238 derivative for expression of LDT<sub>Go</sub> ΔSP, carrying an N-terminal deletion of 30 residues. Kan<sup>R</sup>. | This study |
| FC3619 | pLDT<sub>Go</sub> SP<sub>YcbB</sub> | pSEVA238 derivative for expression of LDT<sub>Go</sub> SP<sub>YcbB</sub>, in which the 52 first aa have been replaced by the SP (first 100 aa) of YcbB<sub>Ec</sub>. Kan<sup>R</sup>. | This study |
| FC3620 | pLDT<sub>Bcn</sub> | pSEVA238 derivative for expression of LDT<sub>Bcn</sub> from *B. cenocepacia*. Kan<sup>R</sup>. | This study |
| FC3621 | pPBP7<sub>Go</sub> | pSEVA238 derivative for expression of PBP7<sub>Go</sub>. Kan<sup>R</sup>. | This study |
| FC3622 | pPBP7<sub>Go</sub> S56A | pSEVA238 derivative for expression of PBP7<sub>Go</sub> S56A point mutant. Kan<sup>R</sup>. | This study |
| FC3623 | pDacB<sub>Ec</sub> | pSEVA238 derivative for expression of DacB from *E. coli*. Kan<sup>R</sup>. | This study |
| FC3624 | pPbpG<sub>Ec</sub> | pSEVA238 derivative for expression of PbpG from *E. coli*. Kan<sup>R</sup>. | This study |
| FC3625 | pET28b(+) | Expression vector with T7 promoter and terminator flanking MCS, and optional C-terminal 6xHis-tag sequence. Kan<sup>R</sup>. | Novagen |
| FC2037 | pET28-LDT<sub>Go</sub> | pET28b(+) derivative for inducible expression of LDT<sub>Go</sub> from *G. oxydans* in *E. coli* B121. Kan<sup>R</sup>. | This study |
| FC3626 | pET28-LDT<sub>Go</sub> C264A | pET28b(+) derivative for inducible expression of LDT<sub>Go</sub> C264 point mutant from *G. oxydans* in *E. coli* B121. Kan<sup>R</sup>. | This study |
| FC3627 | pET28-LDT<sub>Ap</sub> | pET28b(+) derivative for inducible expression of LDT<sub>Go</sub> from *A. pasteurianus* in *E. coli* B121. Kan<sup>R</sup>. | This study |
| FC3628 | pET28-LDT<sub>Bcn</sub> | pET28b(+) derivative for inducible expression of LDT<sub>Go</sub> from *B. cenocepacia* in *E. coli* Kan<sup>R</sup>. | This study |
| FC3629 | pET28-LDT<sub>Bcn</sub> C354A | pET28b(+) derivative for inducible expression of LDT<sub>Go</sub> C354A point mutant from *B. cenocepacia* in *E. coli* B121. Kan<sup>R</sup>. | This study |
| FC2025 | pET28-ldcA | pET28b(+) derivative for purification of LdcA in *E. coli* B121. The protein carries a C-terminal 6-His tag. Kan<sup>R</sup>. | 24 |
| FC1168 | pET28-ldtA | pET28b(+) derivative for purification of LdtA from *V. cholerae* in *E. coli* B121. The protein has a N-terminal deletion of the signal peptide and carries a C-terminal 6-His tag. Kan<sup>R</sup>. | 9 |
| FC2055 | pET22b(+) | Expression vector with T7 promoter and terminator flanking MCS, *pelB* leader sequence for potential periplasmic localization and optional C-terminal 6xHis-tag sequence. Amp<sup>R</sup>. | Novagen |
| FC3630 | pET22-PelB-LDT<sub>Go</sub> | pET22b(+) derivative for inducible expression and purification of PelB-LDT<sub>Go</sub> *G. oxydans* in *E. coli* B121. The first 52 aa of LDT<sub>Go</sub> are replaced by the PelB sequence for expression in the periplasmic space; the protein carries a C-terminal 6-His tag. Amp<sup>R</sup>. | This study |
| FC2778 | pET22-KP27 | pET22b(+) derivative for purification of KP27 from *Klebsiella* phages. Amp<sup>R</sup>. | 23 |
| FC2048 | pSEVA235 | *lacZ* transcription reporter plasmid. Kan<sup>R</sup>. | 73 |
| FC3631 | pSEVA235-Pldt<sub>Go</sub> | pSEVA235 derivative for testing the *ldt<sub>Go</sub>*<br>promoter region (500 bp upstream of the<br>*gox1073* gene). Kan<sup>R</sup>. | This study |

**Table S4.**
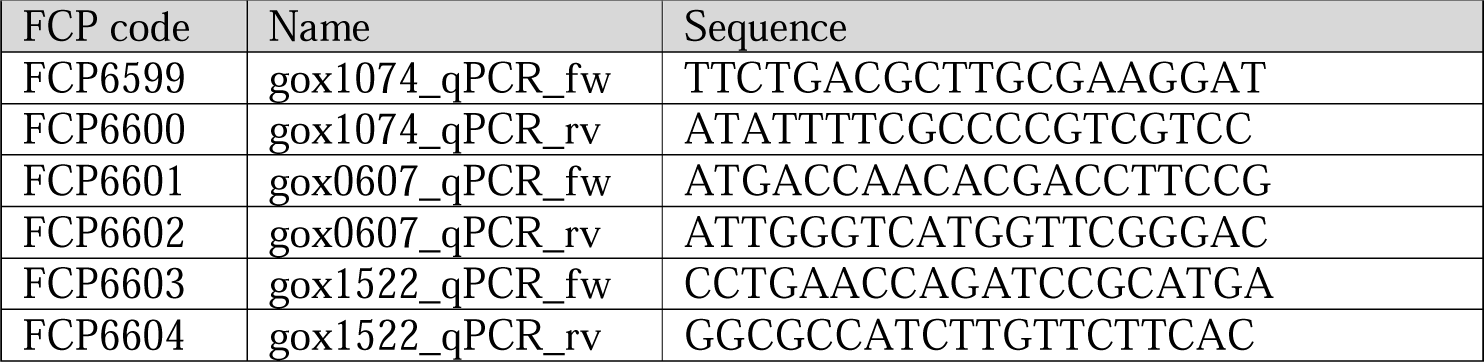
Primers for RT-PCR.

**Table S5.** Identified muropeptides.

| Muuropeptide | Composition | Monoisotopic<br>mass | m/z |  |  |  |
| --- | --- | --- | --- | --- | --- | --- |
|  |  |  | Theoretical | Charge | Observed | Difference |
| Monomers |  |  |  |  |  |  |
| M1 | NAG-NAM-A | 569.243 | 570.251 | 1 | 570.252 | 0.001 |
| M2 (M2<sup>Glu</sup>) | NAG-NAM-A-E | 698.286 | 699.294 | 1 | 699.308 | 0.014 |
| M2<sup>Gln</sup> | NAG-NAM-A-Q | 697.302 | 698.3096 | 1 | 698.313 | 0.003 |
| M2<sup>Phe</sup> | NAG-NAM-A-F | 716.312 | 717.3194 | 1 | 717.314 | -0.005 |
| M2<sup>Trp</sup> | NAG-NAM-A-W | 755.323 | 756.3303 | 1 | 756.330 | 0.000 |
| M3 | NAG-NAM-A-E-mDAP | 870.371 | 871.3784 | 1 | 871.380 | 0.002 |
| M3<sup>Anh</sup> | NAG-NAM<sup>Anh</sup>-A-E-mDAP | 850.344 | 851.352 | 1 | 851.355 | 0.003 |
| M3<sup>NH2</sup> | NAG-NAM-A-E-mDAP<sup>NH2</sup> | 869.387 | 870.394 | 1 | 870.403 | 0.009 |
| M4 | NAG-NAM-A-E-mDAP-A | 941.408 | 942.416 | 1 | 942.432 | 0.016 |
| M4<sup>Anh</sup> | NAG-NAM<sup>Anh</sup>-A-E-mDAP | 921.382 | 922.389 | 1 | 922.389 | 0.000 |
| M4<sup>Met</sup> | NAG-NAM-A-E-mDAP-M | 1001.411 | 1002.419 | 1 | 1002.434 | 0.015 |
| M5 | NAG-NAM-A-E-mDAP-A-A | 1012.445 | 1013.453 | 1 | 1013.456 | 0.003 |
| Dimers |  |  |  |  |  |  |
| D33<sup>Anh</sup> | M3-M3<sup>NH2</sup> (mDAP<sup>3</sup>-mDAP<sup>3</sup>, LD3,3-crosslink) | 1702.704 | 852.360 | 2 | 852.364 | 0.004 |
| D43<sup>NH2</sup> | M4-M3<sup>NH2</sup> (A<sup>4</sup>-mDAP<sup>3</sup>, DD-crosslink) | 1792.784 | 897.400 | 2 | 897.415 | 0.015 |
| D43<sup>NH2 Anh</sup> | M4<sup>Anh</sup>-M3<sup>NH2</sup> (A<sup>4</sup>-mDAP<sup>3</sup>, DD-crosslink) | 1772.758 | 887.387 | 2 | 887.385 | -0.002 |
| D44 | M4-M4 (A<sup>4</sup>-mDAP<sup>3</sup>, DD-crosslink) | 1864.805 | 933.410 | 2 | 933.416 | 0.006 |
| D44<sup>Met</sup> | M4-M4<sup>Met</sup> (A<sup>4</sup>-mDAP<sup>3</sup>, DD-crosslink) | 1924.808 | 963.412 | 2 | 963.411 | -0.001 |
| D44<sup>Anh</sup> | M4-M4<sup>Anh</sup> (A<sup>4</sup>-mDAP<sup>3</sup>, DD-crosslink) | 1844.779 | 923.397 | 2 | 923.402 | 0.005 |
| D45 | M4-M5 (A<sup>4</sup>-mDAP<sup>3</sup>, DD-crosslink) | 1935.842 | 968.929 | 2 | 968.931 | 0.002 |
| D13 | M1-M3 (A<sup>1</sup>-mDAP<sup>3</sup>, LD1,3-crosslink) | 1421.603 | 711.809 | 2 | 711.834 | 0.025 |
| D13<sup>NH2</sup> | M1-M3<sup>NH2</sup> (A<sup>1</sup>-mDAP<sup>3</sup>, LD1,3-crosslink) | 1420.619 | 711.3174 | 2 | 711.316 | -0.001 |
| D14 | M1-M4 (A<sup>1</sup>-mDAP<sup>3</sup>, LD1,3-crosslink) | 1492.640 | 747.328 | 2 | 747.343 | 0.015 |
| D15 | M1-M5 (A<sup>1</sup>-mDAP<sup>3</sup>, LD1,3-crosslink) | 1563.677 | 782.847 | 2 | 782.846 | -0.001 |
| Trimers |  |  |  |  |  |  |
| T444 | M4-M4-M4 (2x mDAP<sup>3</sup>-mDAP<sup>3</sup>, DD-crosslink) | 2788.202 | 930.408 | 3 | 930.410 | 0.002 |
| T144 | M1-M4-M4 (A<sup>1</sup>-mDAP<sup>3</sup>, LD1,3-crosslink and A<sup>4</sup>-mDAP<sup>3</sup>, DD-crosslink) | 2416.037 | 806.354 | 3 | 806.353 | -0.001 |
| T143<sup>NH2</sup> | M1-M4-M3<sup>NH2</sup> (A<sup>1</sup>-mDAP<sup>3</sup>, LD1,3-crosslink and A<sup>4</sup>-mDAP<sup>3</sup>, DD-crosslink) | 2344.016 | 782.347 | 3 | 782.348 | 0.001 |
| Endopeptidase products |  |  |  |  |  |  |
| Tripeptide | E-mDAP-A | 390.175 | 391.183 | 1 | Not found | - |
| Tripeptide<sup>Met</sup> | E-mDAP-M | 450.178 | 451.186 | 1 | 451.189 | 0.003 |
| M43 | M4-tripeptide (A<sup>4</sup>-mDAP<sup>3</sup>, DD-crosslink) | 1313.572 | 657.794 | 1 | 657.795 | 0.001 |
| chain M1-M1 | M1-M1 (β1→4 glycosidic bond) | 1118.460 | 1119.468 | 1 | 1119.463 | -0.005 |
| chain M1-M1-M3 | M1-M1-M3 (2x β1→4 glycosidic bond) | 1968.805 | 985.4101 | 2 | 985.412 | 0.002 |
| chain M1-M1-M4 | M1-M1-M4 (2x β1→4 glycosidic bond) | 2039.842 | 1020.929 | 2 | 1020.928 | -0.001 |

## Supplementary movies

**Movie S1. Molecular dynamics simulations of LDT_Go_.** The video comprises: (i) a rotating view of a rainbow-colored cartoon representation of the X-ray crystal structure, the dots highlighting the gap corresponding to the capping loop that is missing in the electron density map; (ii) a rotating view of our all-atom model of the full-length protein, first displaying all atoms and then only a rainbow-colored cartoon representation; (iii) a molecular dynamics trajectory spanning 350 ns (showing one snapshot every 5 ns) of the LDT_Go_ model that relates to the histogram plot of Fig. S13A, highlighting that motion is largely restricted to the capping loop (C atoms in grey) and parts of the N-terminal Pro-rich belt (C atoms in dark blue); (v) repositioning of the camera to display an alternative orientation (30 frames); (vi) a molecular dynamics trajectory spanning 240 ns (showing one snapshot every 5 ns) of the belt-truncated LDT_Go_ in complex with one peptidoglycan strand (C atoms in violet); and (vi) zoom-in on the active site of LDT_Go_ to show how the catalytic thiol of Cys264 is poised for nucleophilic attack to the carbonyl carbon of L-Ala^1^ (C atoms in pink) of the donor muropeptide (C atoms in violet). The simulations on the apo form of LDT_Go_ were fully unrestrained whereas for the PG complex, (i) C1 and C4 atoms of N-acetylglucosamine and of N-acetylmuramic were restrained with a weak harmonic force constant of 2 kcal mol^−1^ Å^−2^) to preserve the PG lattice architecture, and (ii) a harmonic force constant of 10 kcal mol^−1^ Å^−2^ was employed to keep the Cys264(SG)[L-Ala(C=O) distance between 3.9 and 4.0 Å.

