## Supplementary Figures for "A distinctive family of L,D-transpeptidases catalyzing L-Ala-mDAP crosslinks in Alpha and Betaproteobacteria"

[6] new address: Chr. Hansen A/S, Microbial Physiology, R&D, 2970 Hoersholm, Denmark

Supplementary Figure 1

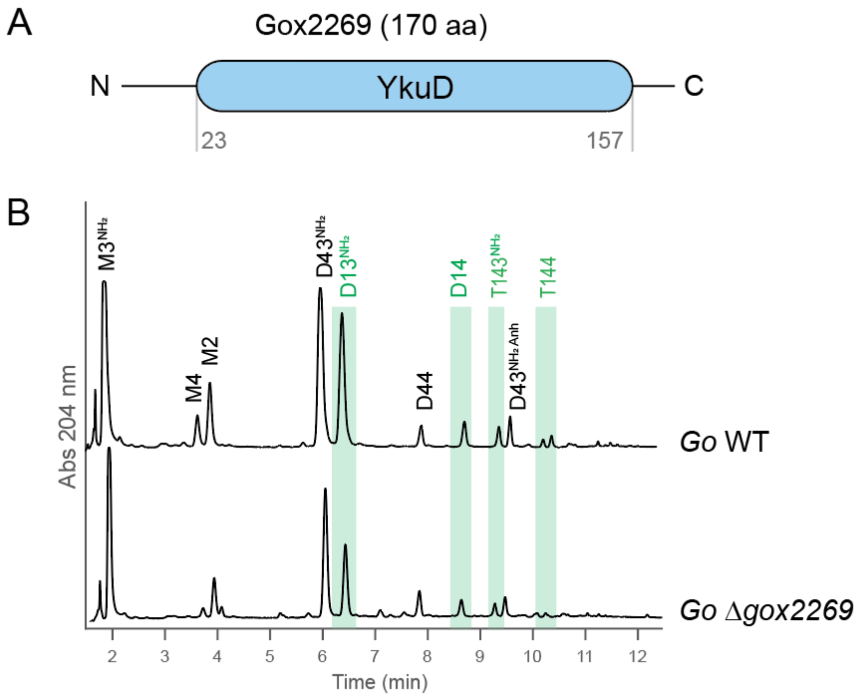

Supplementary Figure 2

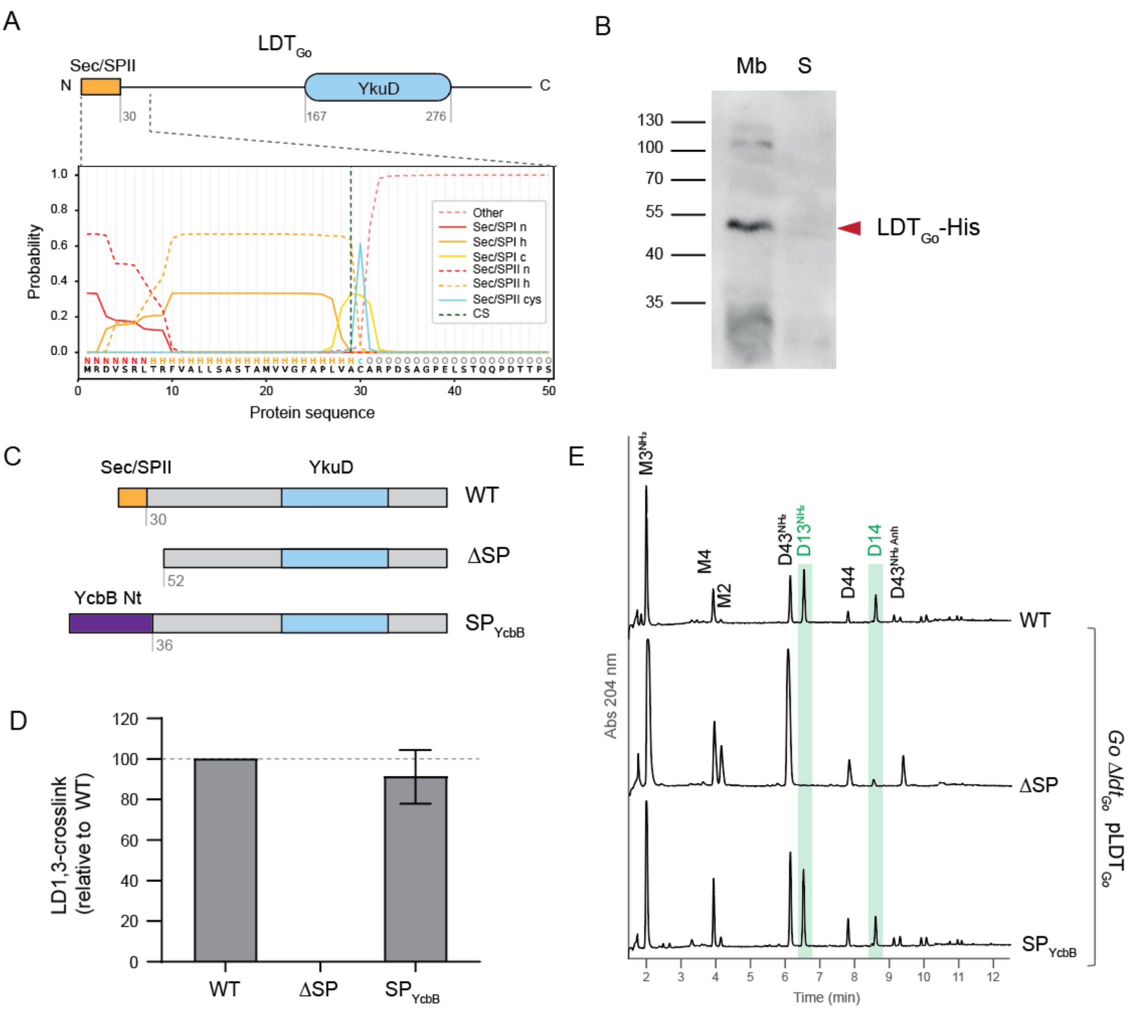

Supplementary Figure 3

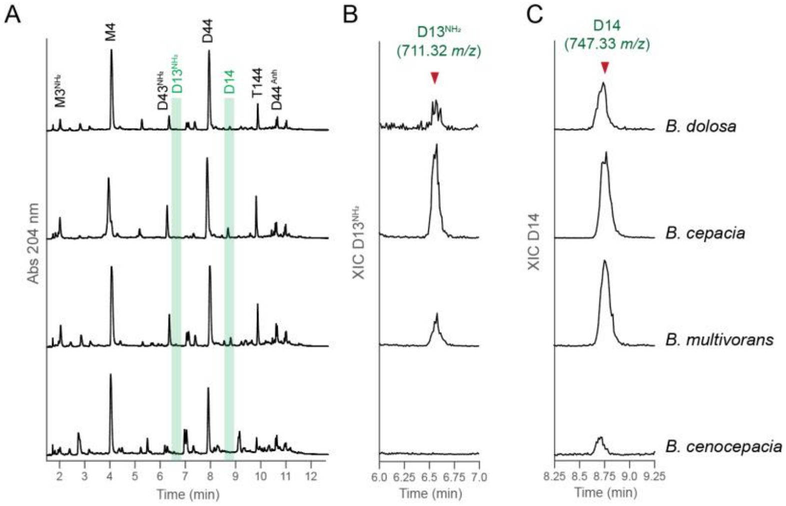

Supplementary Figure 4

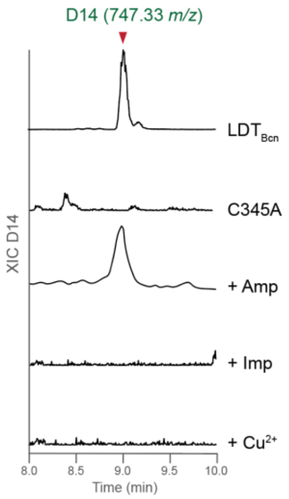

Supplementary Figure 5

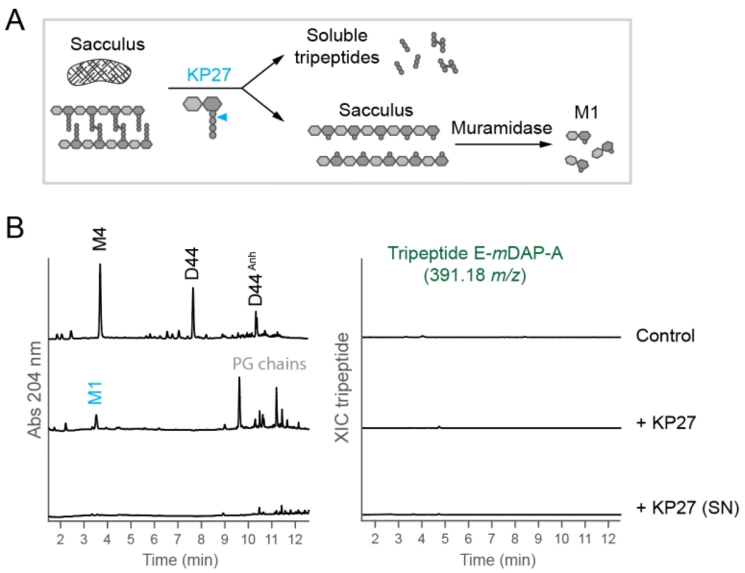

Supplementary Figure 6

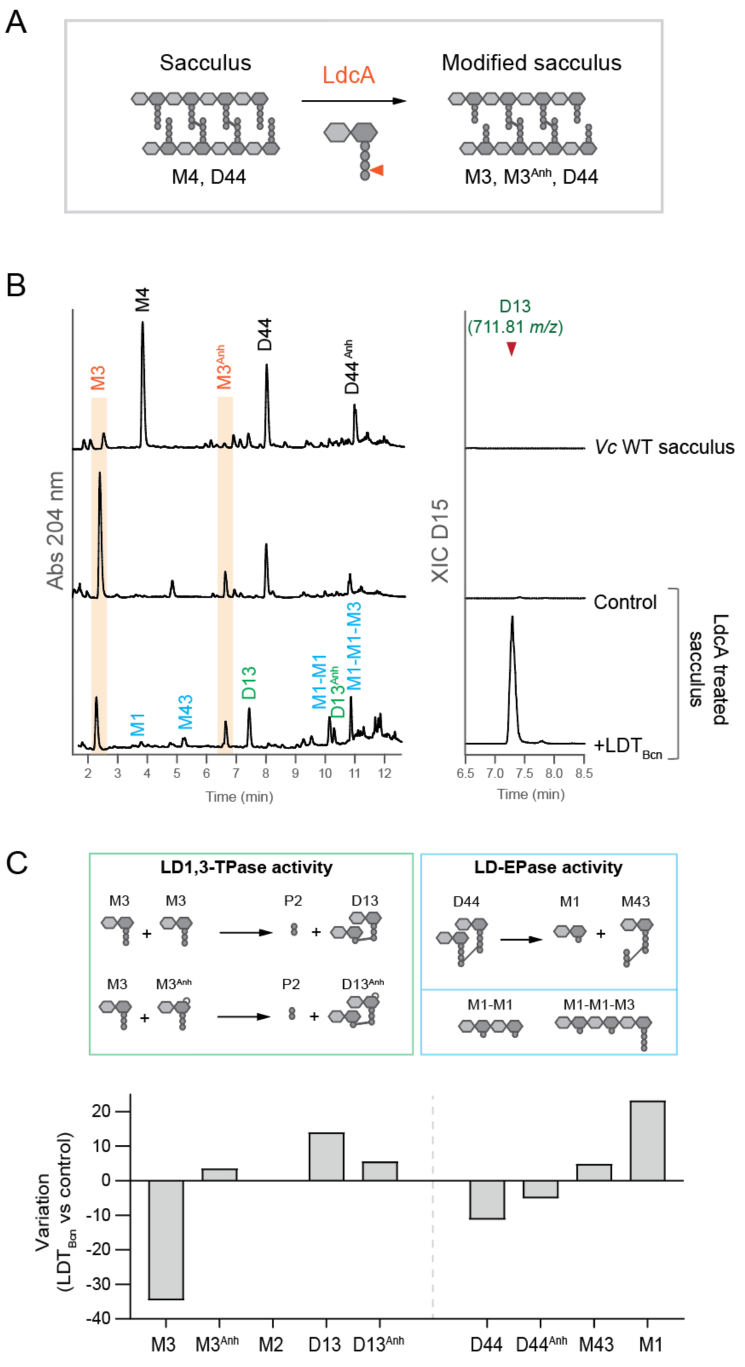

Supplementary Figure 7

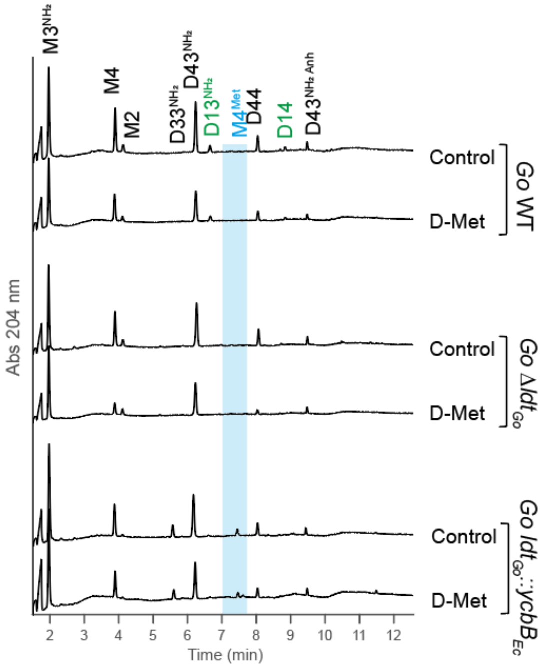

Supplementary Figure 8

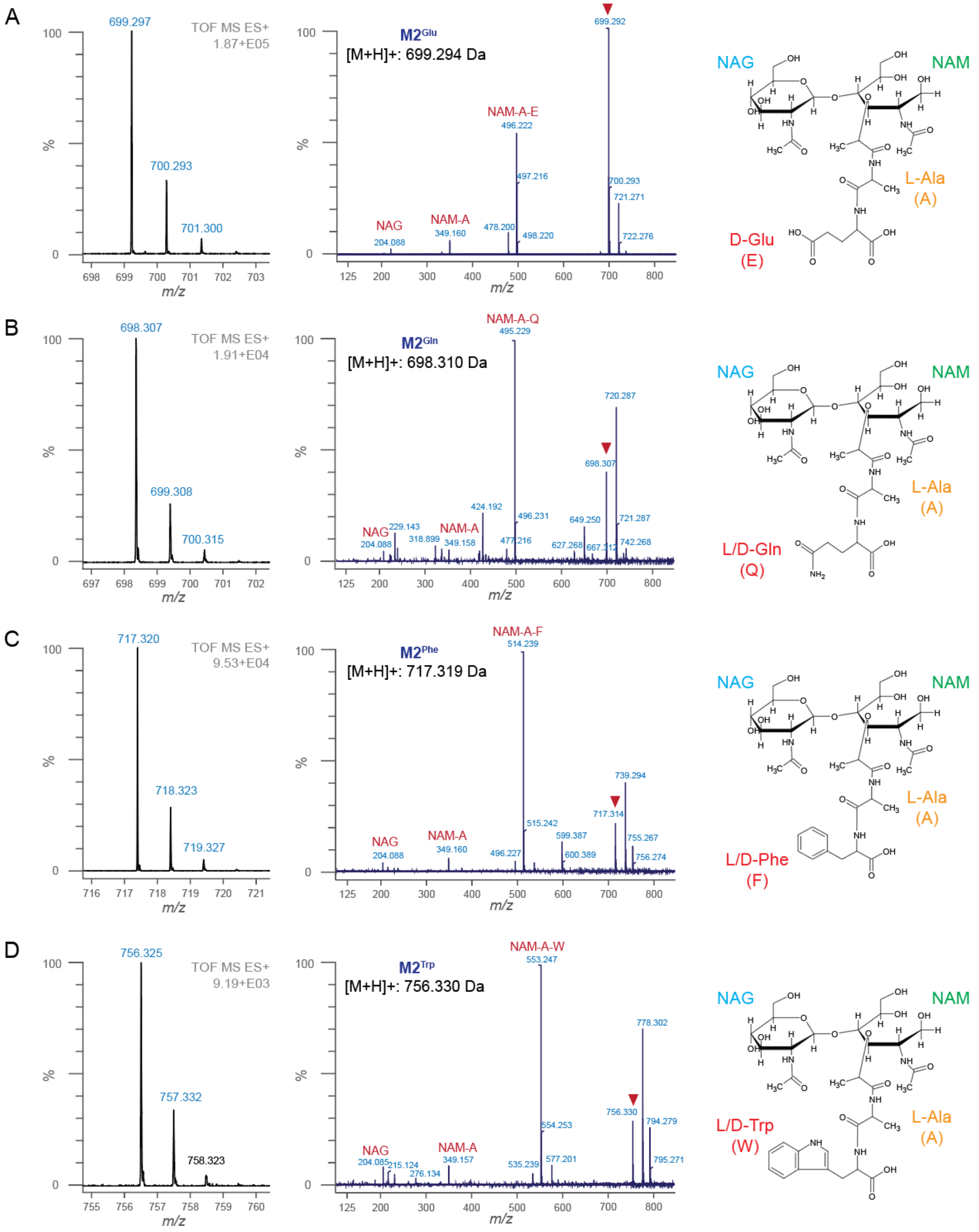

Supplementary Figure 9

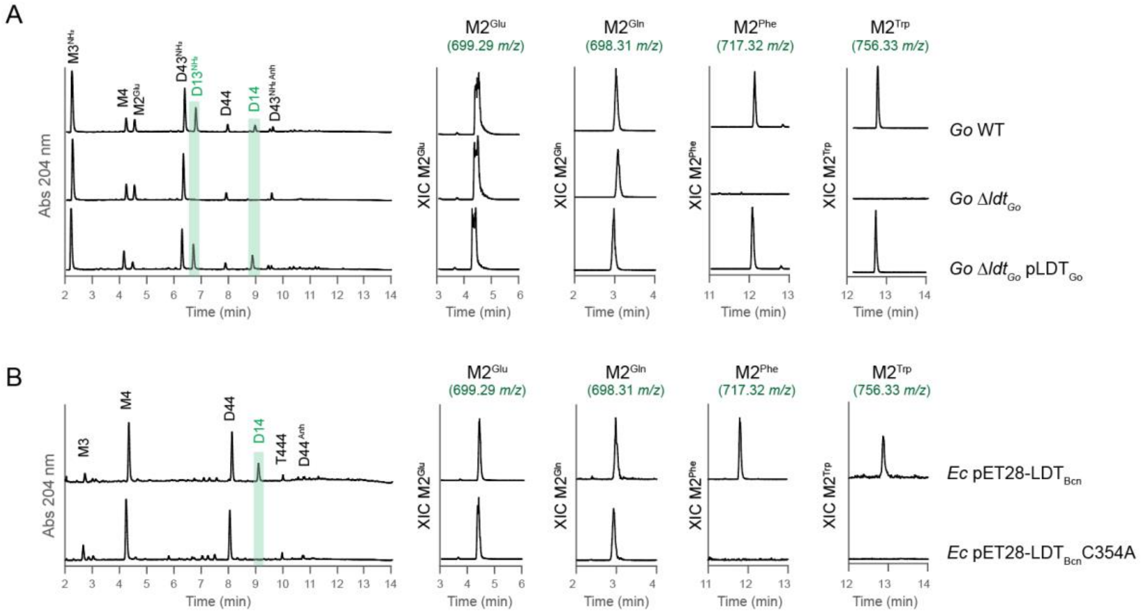

Supplementary Figure 10

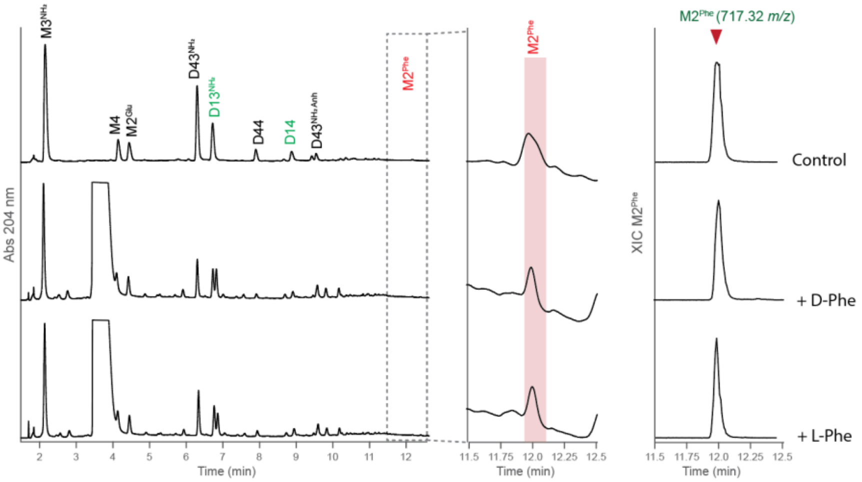

Supplementary Figure 11

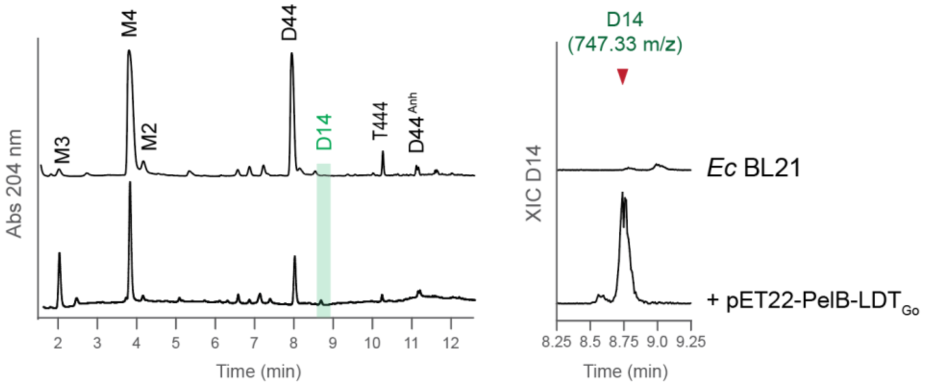

Supplementary Figure 12

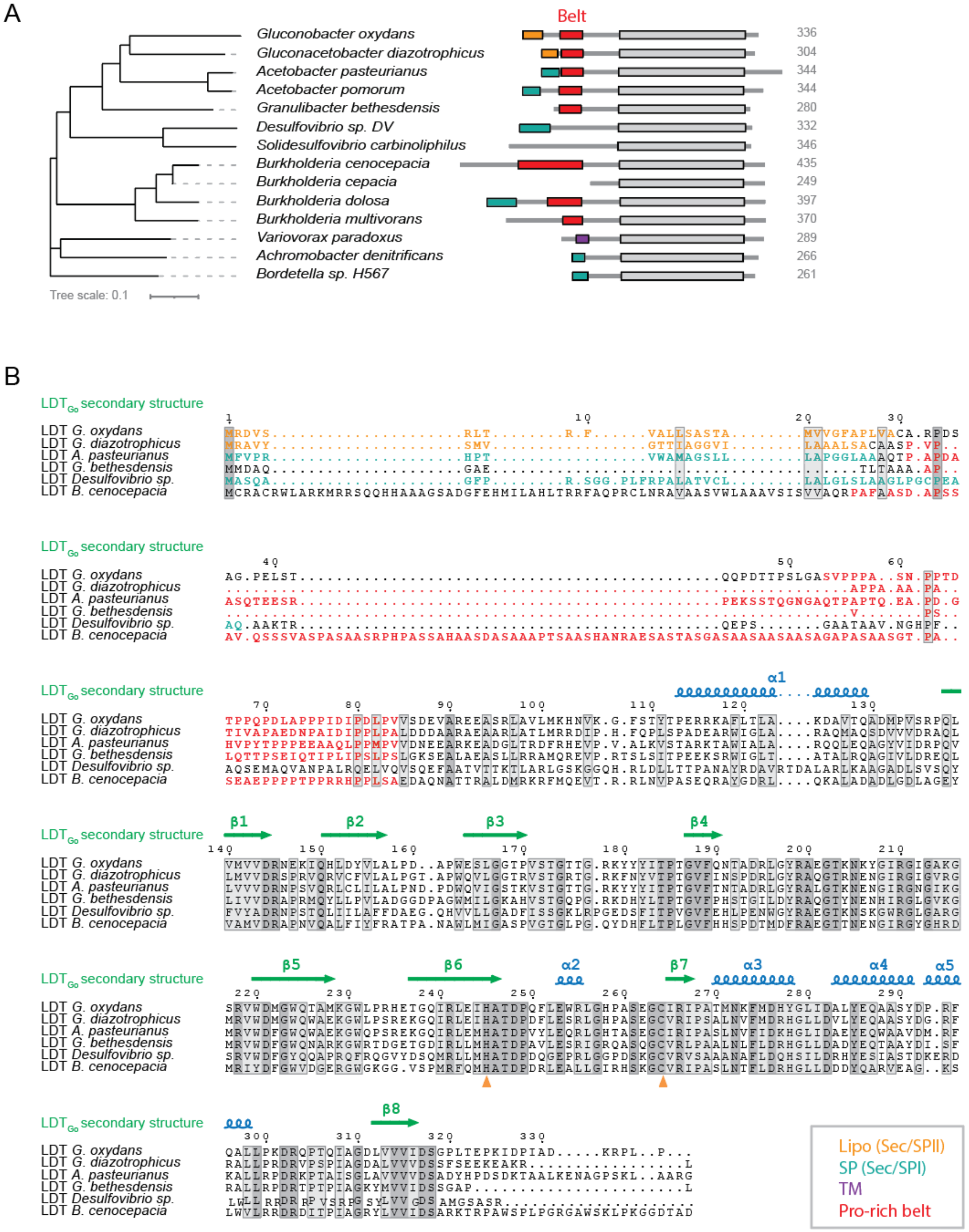

Supplementary Figure 13

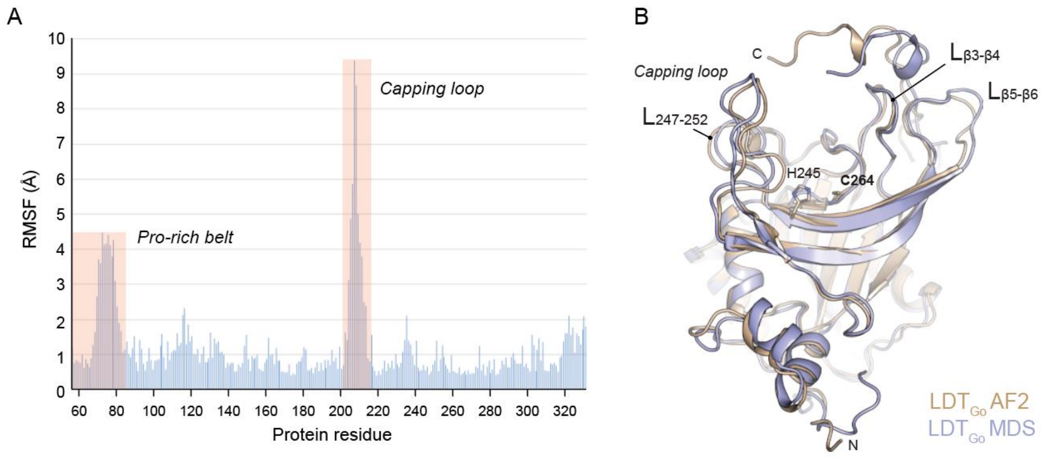

Supplementary Figure 14

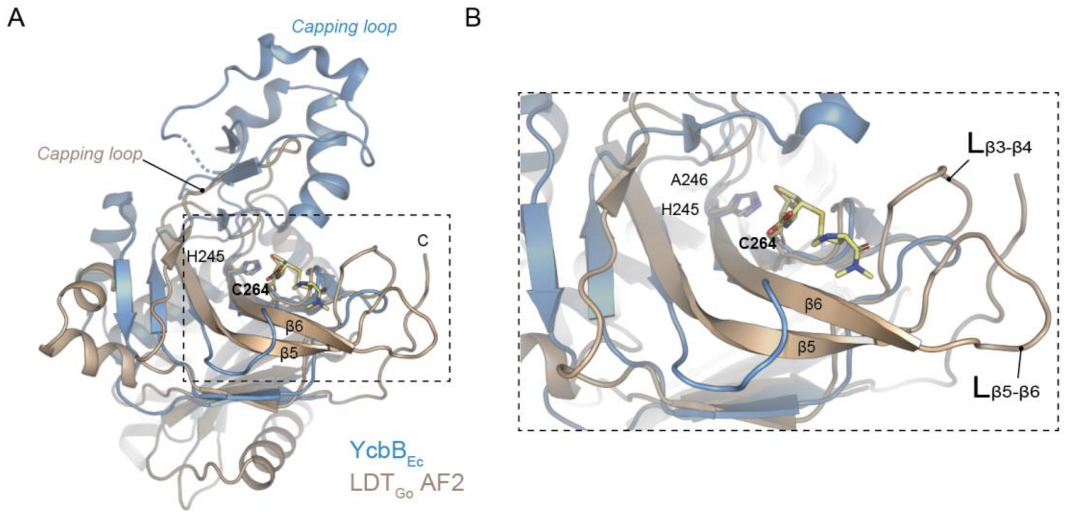

Supplementary Figure 15

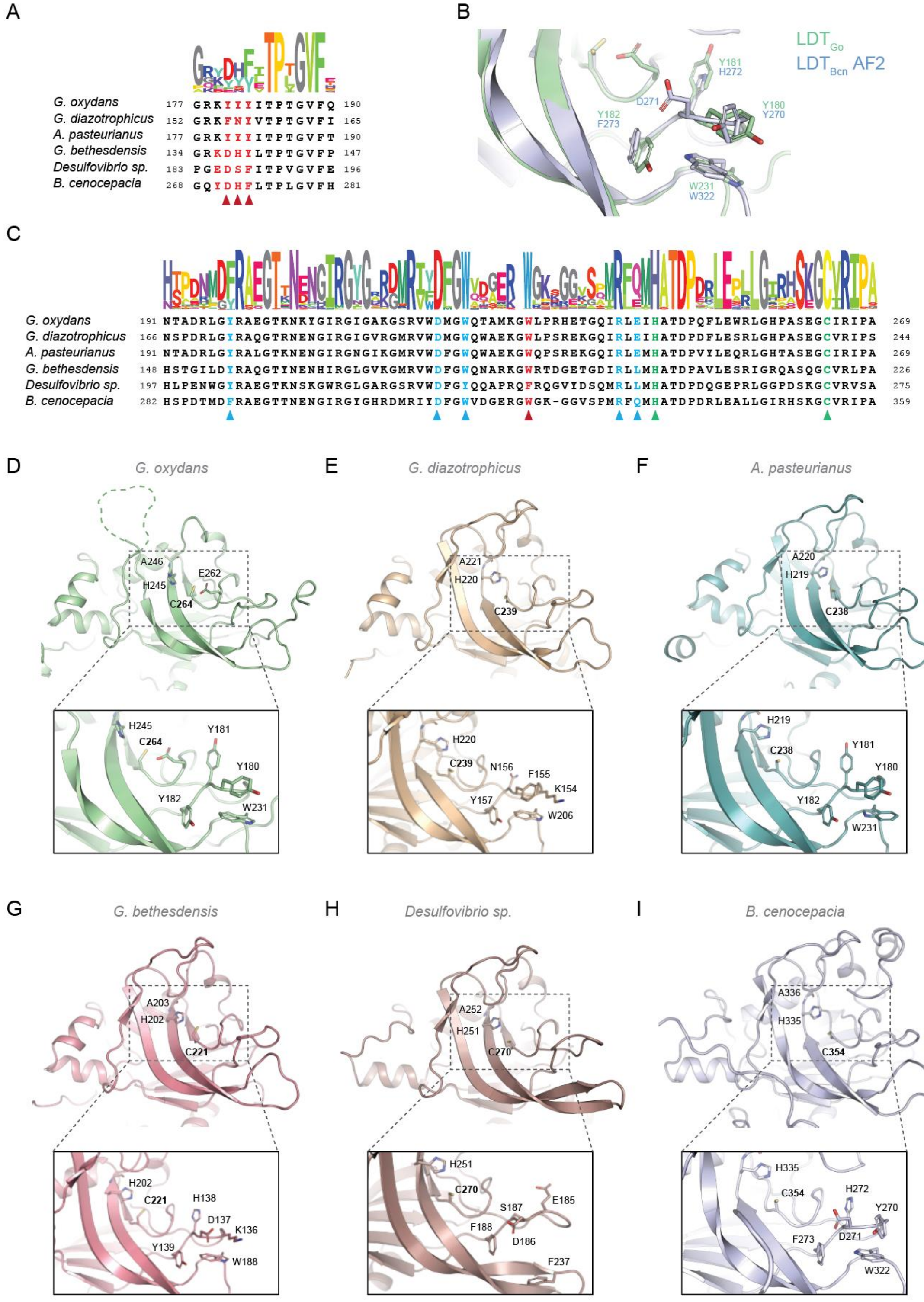

Supplementary Figure 16

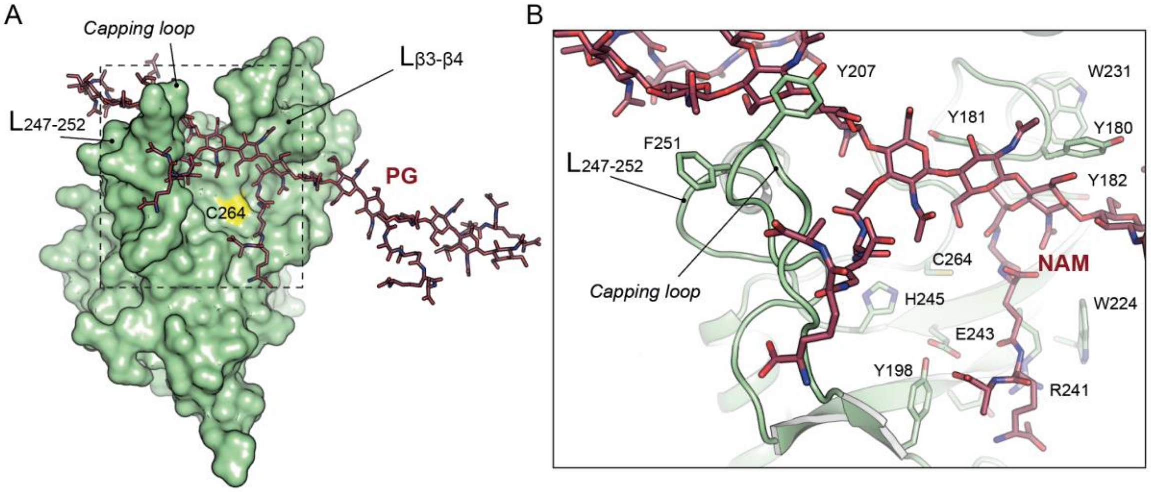

Supplementary Figure 17

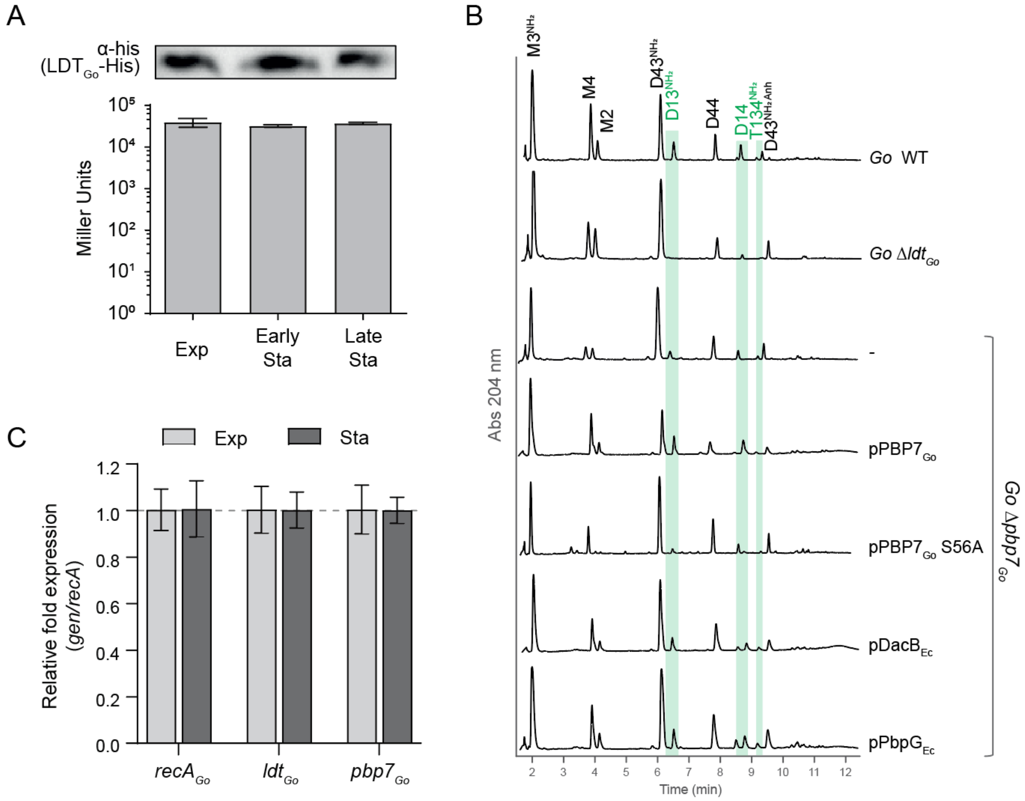

Supplementary Figure 18

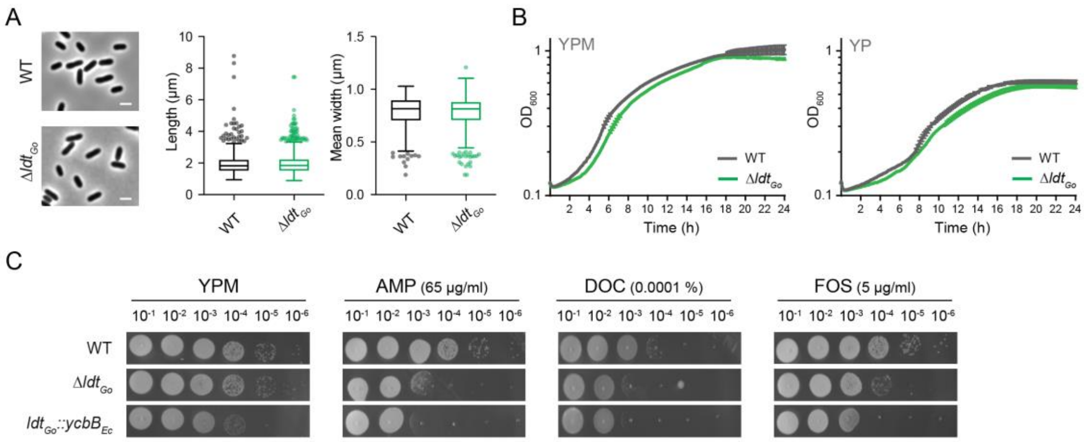

Supplementary Figure 19

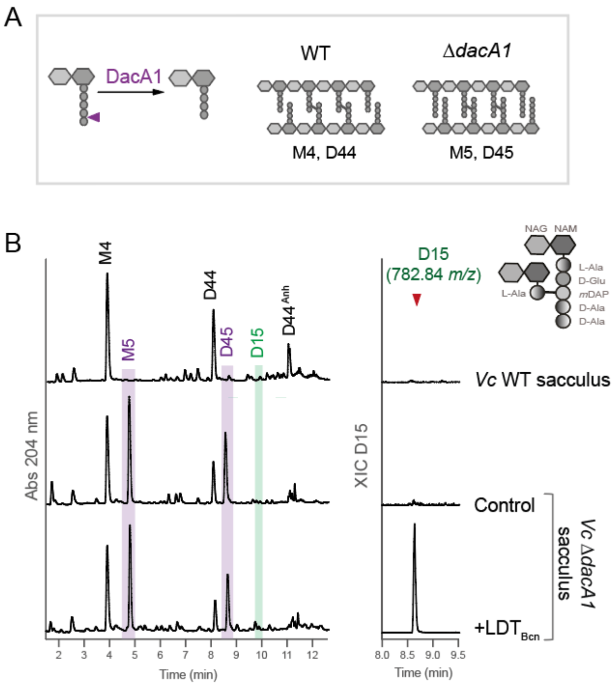

Supplementary Figure 20

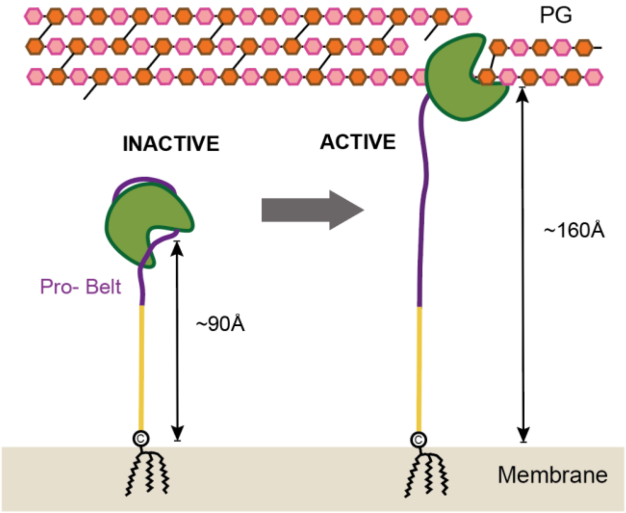
